# All-optical interrogation of excitability during seizure propagation reveals high local inhibition amidst baseline excitability

**DOI:** 10.1101/2023.01.16.524224

**Authors:** Prajay T. Shah, Taufik A. Valiante, Adam M. Packer

**Affiliations:** Krembil Brain Institute, University Health Network, Toronto, ON, Canada; Institute of Biomedical Engineering, University of Toronto, Toronto, ON, Canada; Department of Electrical and Computer Engineering, University of Toronto, Toronto, ON, Canada; Institute of Medical Sciences, University of Toronto, Toronto, ON, Canada; Division of Neurosurgery, Department of Surgery, University of Toronto, Toronto, ON, Canada; Department of Physiology, Anatomy, and Genetics, University of Oxford, Oxford, UK

**Keywords:** Seizures, epilepsy, all-optical, calcium imaging, inhibition

## Abstract

Seizures are classically described as an epiphenomenon of hyperexcitability and hypersynchronicity across brain regions. However, this view is insufficient to explain the complex, dynamic evolution of focal-onset seizures in the brain. Recent studies have proposed mechanisms involving an evolution of excitability driven specifically by a spatiotemporally progressing seizure wavefront. These mechanisms attempt to align the abnormal propagation of neural activity with well-known neurobiological parameters, such as excitation-inhibition balance and neuronal connectivity patterns. We describe a direct test of these mechanisms by performing real-time, *in vivo* investigations of excitability in the acutely epileptic state and during seizure propagation. We used all-optical interrogation to test single-neuronal and local-circuit excitability in the epileptic brain. We demonstrate a surprising paradox during the acutely epileptic state, wherein the brain becomes susceptible to large synchronous inputs, yet single-cell excitability is largely maintained at baseline levels. At a finer scale, excitability of neurons at the single-cell level is related to their distance from the seizure wavefront. Local circuit excitability is increased in the distal penumbra but, crucially, we find inhibition in close proximity to the seizure wavefront. This is in contrast with previously suggested notions of widespread inhibition outside the direct area of action during a focal-onset seizure. These experimental results provide the first direct, *in vivo* evidence for the precise spatial scale over which single-cell excitability dynamics evolve during seizure propagation, providing support for local inhibitory restraint of seizure propagation.

## Introduction

The dynamic activity of neurons, whether physiological or pathological, is determined by a variety of factors which together influence the excitability of those neurons. The most important determinant of the real-time excitability of a given neuron is the inputs it receives and integrates from all of its presynaptic partners within the overall network: Hyperpolarizing (inhibitory) inputs decrease neuronal excitability and depolarizing (excitatory) inputs increase neuronal excitability (1). In the study of brain disease, the characterisation of neuronal excitability can help identify key pathophysiological mechanisms. Epilepsy is the quintessential disease of excitability in the brain, in which the brain becomes susceptible to repeated, generally unprovoked seizures. These seizures, which are the defining disorder of epilepsy, are distinctive episodes of large scale synchronisation and maximally intense neuronal activity. Focal-onset seizures are a sub-class of seizures in which seizure activity begins in a localised (most often cortical) region in the brain (2, 3), and propagates to encompass multiple brain regions. Despite the extreme and disrupted state that the brain reaches during a seizure, seizure propagation must still be fundamentally constrained by wellknown neurophysiological functions. Although a wide array of neurological lesions can lead to seizures, the mechanisms by which a given neuron enters a seizure-state – either at the focal initiation site or through seizure propagation – are more limited (4). The propagation of seizure activity remains the most mysterious and resistant to clinical management strategies (5). How are neurophysiological functions disrupted to lead to the propagation of seizure activity in the epileptic brain?

The propagation of seizures at the micro-scale level in both human and animal models follows two key characteristics: 1) a distinct boundary of intense, tonic firing activity demarcates the “seizure wavefront”, which spatially divides the non-seizing and actively seizing regions of the brain termed “seizure penumbra” and “seizure core”, respectively (6, 7), and 2) the propagation of this seizure wavefront across a population of neurons is highly reliable, paralleling the generally stereotypical nature of seizure propagation patterns recorded in epileptic patients. A striking feature of seizure propagation is the paradoxical disparity in the intense firing of neurons inside the seizure and the low-firing state outside the seizure in the penumbra region, known as “surround inhibition” (6, 8, 9). It is hypothesised that this occurs due to a high activation of feed-forward inhibition in the penumbra driven by the seizure core leading to reduced neuronal excitability and firing in the penumbra (6, 10–12); and the eventual breakdown of this inhibition enables the propagation of seizure to evolve forward (13–16). This was most recently formalised in a computational 2-D neuronal network model, where the seizure wavefront drives cell-type specific evolution of microscale neural network activity dynamics, and strongly affects neuronal excitability in front of itself by causing a complete collapse of feedforward-inhibition, a subsequent steep rise in excitability and thereby becomes the primary driver of its own propagation (17).

The steep rise of excitability that pushes a given neuron into a seizure state is governed by multiple mechanisms, such as raised extracellular [K^+^] and a usage-dependent collapse of GABAergic inhibition (via a depolarized *E*_*GABA*_) to oppose incoming excitatory inputs (17, 18). It is proposed that these take place within a neuronal population directly in front of the seizure wavefront. However, the combination of depolarized *E*_*GABA*_ and electrical stimulation is not sufficient to trigger ictal events, but can act as an adjunct to a primary chemoconvulsant (19). However, inhibition maintains an anti-ictal effect at the seizure onset site for up to 2secs after seizure onset (20). Inhibition has been found to both trigger, enhance or interrupt epileptic activity in both brain slices and *in vivo* (20–23). Thus, the mechanism and role of altered inhibition in seizure remains to be fully elucidated. Specifically, it remains to be determined how inhibition functions in relation to the seizure wavefront, which is the critical site where neurons are transitioning from non-seizing to seizing states.

Many studies of the role of inhibition in regulating neuronal excitability during seizure propagation have been restricted to reduced settings such as brain slices or computational models (but see also (6, 8, 9)) and Liou et al., (2018) who describe surround inhibition *in vivo*). However, a population of neurons *in vivo* additionally receives macro-scale influences such as thalamic, cortico-cortical and sub-cerebral input that contribute to their dynamically-evolving behaviour (24–26). During the event of a seizure, it is unclear how the proposed micro-circuit mechanism of propagation relates to macro-scale circuit influences on neuronal excitability (27). Indeed, there is increasing evidence that macro-scale influences play a role in epilepsy and seizure propagation. Focal-onset epilepsy has been found to exhibit lasting hyperexcitable networks across the brain that may be viable therapeutic targets during seizure propagation (28–32). Large field-of-view widefield calcium imaging shows that epileptic activity propagation over large distances respects the functional, topological connectivity structure of cortex And, it is proposed that the activation of inhibition in distant regions maintains a low-firing state of penumbral neuronal populations that are characterised by large-amplitude EEG signals (6, 10). Furthermore, subcortical routes and neuromodulator influences have also been implicated during a seizure. These *in vivo* studies, using both fMRI and electrophysiology measurements, have found a deactivation of remote regions during focal limbic seizures. These have been suggested to be related to a network effect of the activation of inhibitory circuits in the hypothalamus and the lateral septum caused by decreased cholinergic activity in the cortex in the duration of the seizure itself (33). These studies highlight that the mechanisms dictating the propagation of seizures may involve large-scale brain networks and their neurophysiological influences on excitability which are unique to the *in vivo* condition, and thus need to be studied in this setting.

A key feature of the *in vivo* condition of neurons is that dynamical neuronal excitability is additionally influenced by the conductance state (set by the total number of active synaptic inputs) of post-synaptic neurons. Thus, a deeper understanding of the neuronal processes underway during a seizure can be achieved by directly probing the excitability of neurons located in the penumbra of a seizure *in vivo*. This parallels the measurement of subthreshold dynamics in neurons in the *in vivo* and awake setting which has revealed critical features unique to the behaving state of the brain (1, 34). These studies have highlighted the significant role of shunting inhibition caused by the influence of many-fold greater background synaptic currents present in the *in vivo* state, as compared to the *in vitro* setting (35–37). This notably affects the gain of the close relationship between excitability and neuronal firing (38). Specifically, a low conductance state with relatively little synaptic input raises the excitability of neurons, whereas a high conductance state with greater synaptic inputs lowers the excitability of neurons. Given the intensely heightened scale of neuronal activity during a seizure, it is likely that increased activity across many brain networks impact neuronal excitability during the evolution of seizure, in a manner that is unique to the *in vivo* condition.

The measurement of neuronal excitability requires the simultaneous delivery of input and measurement of neuronal output. Traditionally, this has required intra-cellular access, making it exceptionally difficult to measure single-neuronal excitability across multiple individual neurons *in vivo*. Instead, a highly-targeted and scalable approach for measuring neuronal excitability would be to apply all-optical interrogation, which combines two-photon optogenetic photostimulation and two-photon calcium imaging (39). This approach can in principle measure the excitability of all optically-accessible neurons at single-neuron resolution in the *in vivo*, awake, behaving and minimally-invasive setting. For example, all-optical interrogation found increased excitability in “non-place” CA1 neurons (40).

To directly assess the excitatory- and inhibitory-mechanisms at play in epilepsy and seizure propagation, we leverage all-optical interrogation to measure single-neuronal excitability in the awake brain *in vivo*. We demonstrate that during seizure propagation, there is an increase in the excitability of neurons which are distal to the seizure wavefront, while proximal neurons show a striking decrease in excitability. This occurs while single-neuron excitability is maintained at baseline levels in the epileptic (interictal) state. Contrary to previous suggestions, these findings suggest a previously unappreciated complex distribution of spatial excitability and provide fundamental insights into the cellular mechanisms at play during seizure.

## Results

We performed two-photon calcium imaging and optogenetic photostimulation experiments of acute, focal-onset seizures in head-fixed mice (Figure 1A). We utilised the focal injection of 4-aminopyridine (4-AP), a K^+^ channel blocker which induces epileptiform activity and local neuronal hyperexcitability by lowering action potential threshold and increasing presynaptic excitatory neurotransmission (41), to generate an acute model of focal-onset epilepsy in the brain, similar to previous calcium imaging studies of focal-onset seizure (42, 43). Briefly, on each experimental day, the experimental field-of-view (FOV) was chosen near the centre of the cranial window based on optimal viral expression at a depth of 150-250 µm below the pia to target layer 2/3. This experimental FOV was >1mm away from the site of the focal 4-AP injection where a local field potential (LFP) recording pipette was also inserted in parallel. The experimental FOV and pipettes remained fixed for the entire duration of each experiment. Overall, this preparation allowed for an induction of an acutely epileptic state with focal seizure onset, LFP electrophysiological recording at the site of focal 4-AP injection and all-optical interrogation distal to the 4-AP ictal core.

**Figure 1:**
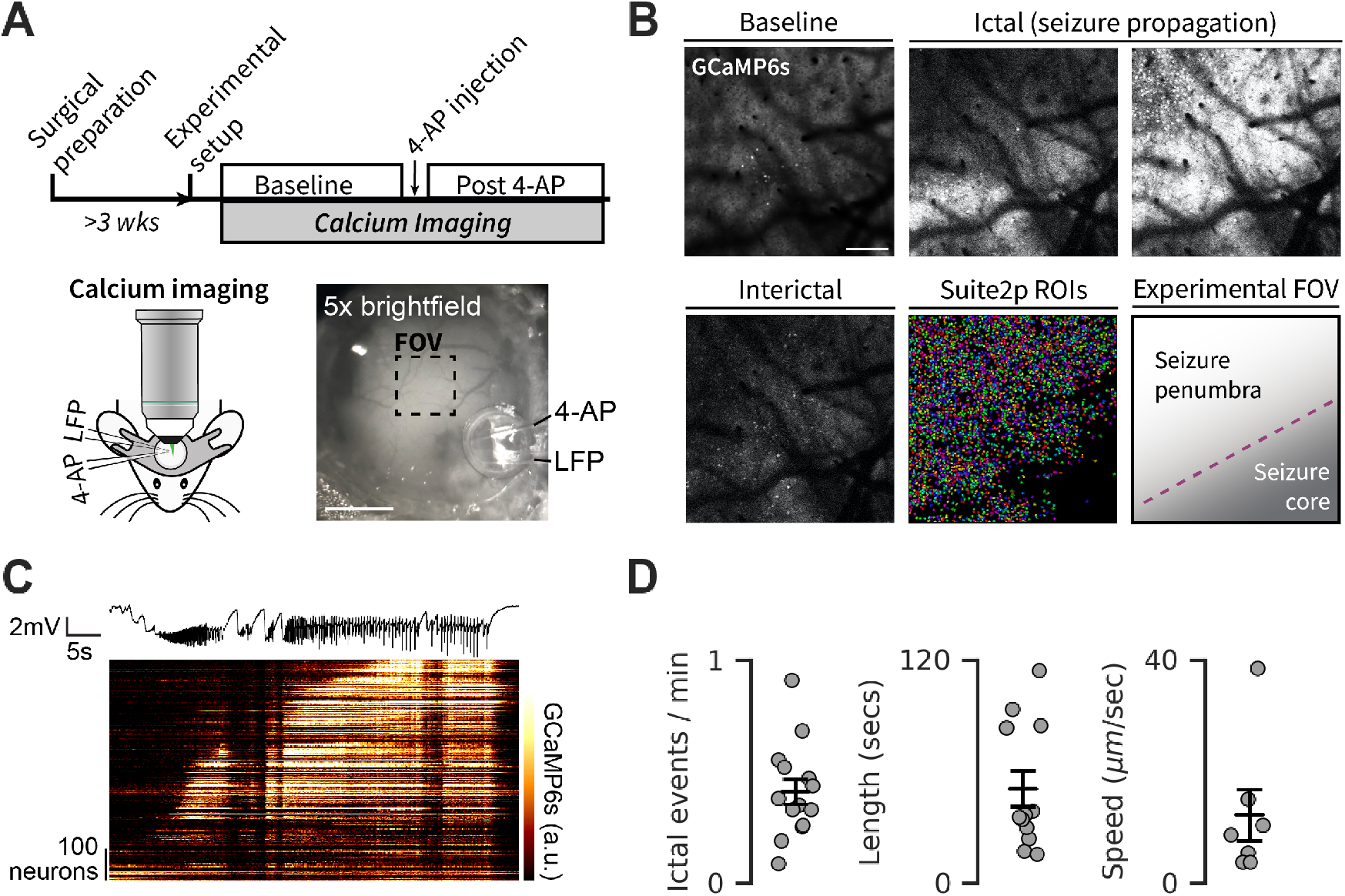
Two-photon neuronal calcium imaging of acute, focal-onset seizures in cortex A. Experimental workflow (top) and setup of simultaneous in vivo two-photon imaging, local field potential (LFP) recording and 4-aminopyridine (4-AP) injection in awake, head-fixed mice over somatosensory cortex. Calcium imaging was peRF-01ormed in CamkIIa-expressing excitatory neurons transgenically or virally induced to express GCaMP6s, and virally induced C1V1-Kv2.1-mScarlet for optogenetic experiments. 4-aminopyridine (4-AP) was injected near the site of LFP recording and > 1mm away from the experimental microscope field of view. Bottom right: Brightfield image of cranial window showing locations of imaging FOV, 4-AP injection pipette and LFP-recording pipette during a representative experimental session. Scale bar = 1 mm B.Imaging experiments were peRF-01ormed at the same microscope FOV during baseline (pre-4-AP injection), interictal (post-4-AP injection, no seizure), and ictal (post-4-AP injection, during seizure). Shown are multi-frame average projections of a representative calcium imaging time-series during an experiment. (Bottom middle) Cell masks collected from Suite2p processing of baseline calcium imaging. (Bottom right) During seizure propagation the seizure wavefront divides the seizure core from the seizure penumbra. Scale bar = 200 µm. C.Calcium imaging signal of neurons from imaging FOV (bottom, heatmap) and corresponding LFP (top, black trace) during one representative seizure. D.Average frequency of seizure incidence (left), average duration of seizures (middle) and average propagation speed of seizure after 4-AP injection measured from calcium imaging experiments.

### Calcium imaging in a model of acute epilepsy

Focal 4-AP injection in the brain induces an acutely epileptic state in which repeated, spontaneously occurring focal-onset seizures arise. We used the simultaneously collected LFP signal, which provided a high temporal resolution electrophysiological signal from the seizure onset site, to mark the onset and offset of seizures during all experiments. The 4-AP injection remains local as seen by co-diluted fluorescent RhodB, but generates seizures which can propagate throughout the brain, as recorded using widefield calcium fluorescence imaging across both hemispheres (Suppl. Figure 1). Simultaneous LFP recording in the contralateral hemisphere confirms this widespread propagation (Suppl. Figure 1, (10, 44).

Widefield calcium imaging provides greater spatial information about neuronal activity than the LFP signal. The widefield calcium fluorescence signal also correlates highly with the LFP signal from the same location (Suppl. Figure 2). This allows the widefield calcium fluorescence signal to be used as a proxy for the LFP at a given region. Additionally, the average widefield calcium fluorescence signal at the 4-AP injection site is heightened even during interictal periods, the widefield calcium fluorescence signal of regions outside the injection site remains low during interictal periods before being heightened during ictal events (Suppl. Figure 1). Finally, for a brief moment following seizure termination, the calcium fluorescence signal is drastically and synchronously suppressed across all regions (Suppl. Figure 1). Both the widefield calcium and LFP signals likely reflect total post-synaptic currents rather than neuronal spiking activity (45, 46). In summary, these data in aggregate show that focal 4-AP injection leads to locally increased activity, but this does not lead to increased activity at distal locations until the seizure initiates.

In contrast to widefield imaging, two-photon imaging provides single-cell resolution imaging of thousands of neurons, here during baseline, interictal and seizure states (Figure 1B). Two-photon calcium imaging shows that during a seizure, there is a protracted progression of the seizure across space which passes through the experimental FOV. During this process, there is a distinct area of intense activity that corresponds to the propagating seizure wavefront (Figure 1B, see also Wenzel et al., 2017). Two-photon imaging allows seizure propagation to be directly observed in the imaged population of neurons as a gradual recruitment of neurons into seizure (ordered by time delay to reach 75% of the maximum signal, Figure 1D). Overall, this experimental paradigm allows for direct visualisation of seizure propagation across a large neuronal population and raises questions about the processes that lead to these characteristics of seizure propagation.

### Activity of interneurons during propagation of focal 4-AP seizures

The regulation of excitability in the brain is achieved by a tight balance of inhibitory and excitatory inputs (47, 48). In a seizure, Liou et al., (2018) have previously found that focal seizure onset is associated with a fast rise in the activity of inhibitory neurons in the ictal penumbra. However, the activity of excitatory neurons was not simultaneously measured, so it remains unclear whether the increased activity of inhibitory neurons distal to the seizure onset site may regulate excitatory activity. We performed cell-type specific two-photon calcium imaging of inhibitory and excitatory neurons in Nkx2.1-Cre:::mCherry mice virally injected to express hSyn-GCaMP7f in all neurons in the same experimental preparation described above (Suppl. Figure 3A). This preparation allowed us to perform calcium imaging of 52 +/-13 (mean +/-stdev.) Nkx2.1^+ve^ inhibitory neurons from each animal (N = 3 animals).

Interestingly, we found that following seizure onset, the rise in GCaMP7f signal Nkx2.1^+ve^ inhibitory neurons in the ictal penumbra was also closely associated with Nkx2.1^-ve^ neurons and the overall GCaMP7f signal imaged at the same FOV (Suppl. Figure 3B). Indeed, the recruitment of Nkx2.1^+ve^ cells into propagating seizures was not significantly different from their local surrounding region (measured as the average of a 100µm annulus around each respective Nkx2.1^+ve^ cell) across a diverse range of individual seizure propagation speeds (Suppl. Figure 3C). Thus we do not find any evidence of a divergence of activity patterns between excitatory neurons and inhibitory neurons during the propagation of a focal-seizure in vivo.

### Neuronal population excitability in acute epilepsy

Our primary goal in this study is to study how neuronal excitability is affected outside of the seizure onset zone in a model of focal-epilepsy? We first tested neuronal population excitability in the acutely epileptic state using combined calcium imaging and widefield one-photon optogenetic photostimulation *in vivo*. This type of stimulation indiscriminately activates a large number of opsin-expressing neurons.

In mice co-expressing the somatically-targeted opsin C1V1-Kv2.1 and GCaMP6s in layer 2/3 of somatosensory cortex (Figure 2A), we performed widefield optogenetic photostimulation, and measured the photostimulation evoked signal with calcium imaging (quantified as the average of the entire experimental FOV) and in the simultaneously recorded LFP. We found a robust and reliable response to photostimulation under baseline conditions (before 4-AP injection) in both the population GCaMP fluorescence signal (averaged across the FOV) as well as the LFP signal (Figure 2B). Note that the central location of viral opsin injection within the cranial window constrained direct (i.e. optogenetic) stimulation to C1V1-expressing neuronal somata which were located greater than 1mm away from the 4-AP injection and LFP recording site. Thus, the average calcium fluorescence signal response represents the response of directly photostimulated cells, whereas the LFP signal likely represents the downstream synaptic activity arising from the directly photostimulated cells since the LFP recording site is outside of the C1V1-opsin expression area.

**Figure 2:**
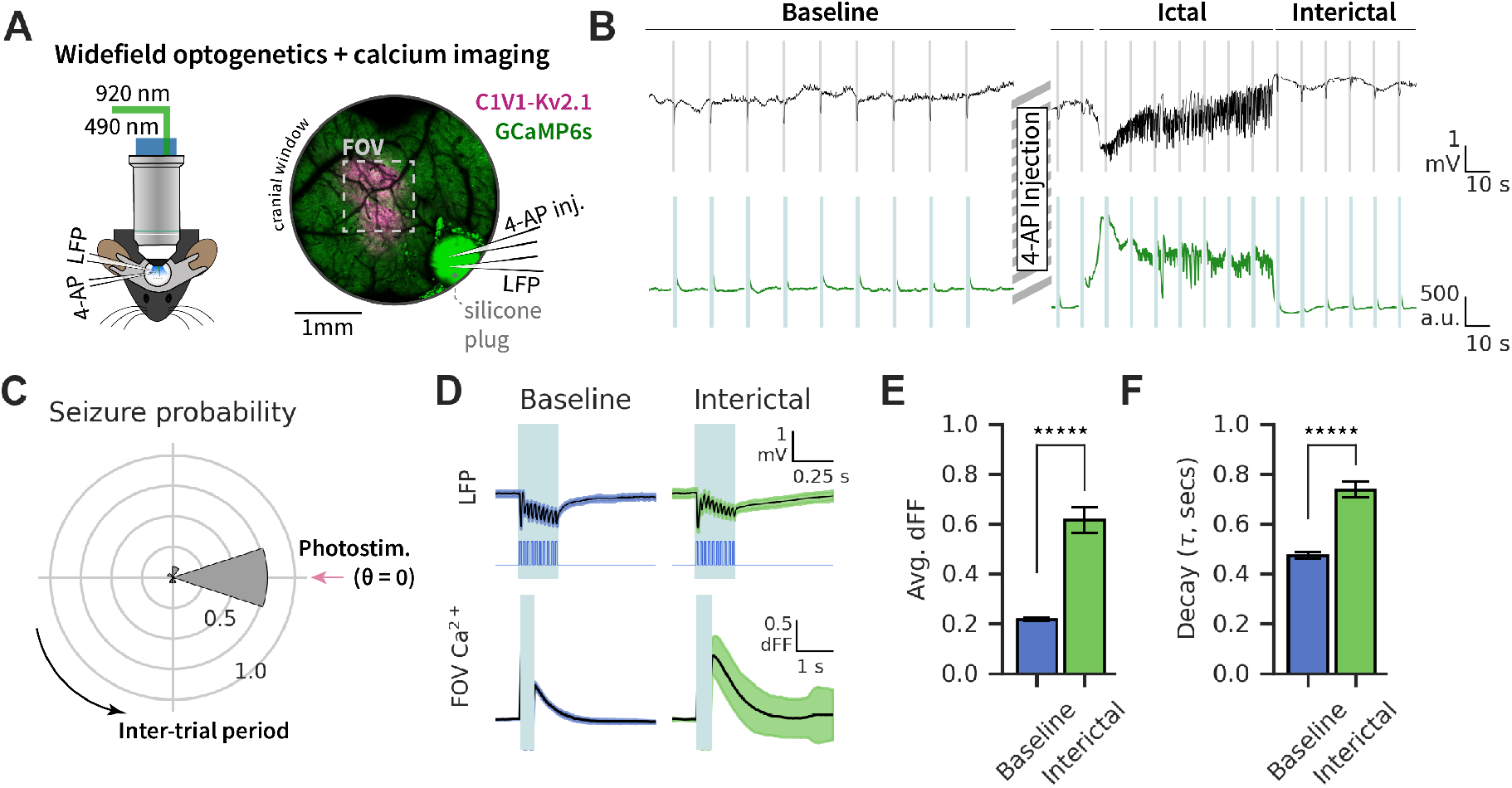
Widefield one-photon stimulation and calcium imaging in the acutely epileptic state A.Left: Experimental setup for combined head-fixed two-photon calcium imaging and widefield optogenetic photostimulation with local-field potential (LFP) recording and 4-aminopyridine (4-AP) injection. Right: Two-photon image of representative cranial window showing viral co-expression of GCaMP6s (green) and C1V1-Kv2.1-mScarlet (magenta). 4-AP and LFP pipettes are inserted through the access hole in the cranial window after removal of the silicone plug. B.LFP recording (top) and corresponding mean FOV fluorescence calcium imaging signal (bottom) from a single representative experiment before and after 4-AP injection. Blue stripes represent widefield photostimulation trials. Highspeed shutters covering the imaging detectors were triggered for protection during widefield illumination. C.Probability of seizure occurrence relative to photostimulation timing across all seizure events (1 sec bins, N = 5 mice, p(V test for circular non-uniformity) = 1.3e-4). D.Photostimulation-timed average response (+/-standard dev.) of LFP signal and widefield illumination TTL trigger output (top), and FOV-average calcium imaging signal (bottom) from a representative experiment. Blue span represents widefield illumination. E.Average photostimulation response magnitude during baseline and interictal states (N = 5-6 mice per group, 360 baseline trials and 172 interictal trials (trials inducing seizures were excluded); t-test: *****p<1e-24). F.Average decay constant of photostimulation response across baseline and interictal states (N = 5-6 mice per group, 359 baseline trials and 156 interictal trials (trials inducing seizures were excluded), t-test: *****p<1e-17). Decay constant calculated as the time (secs) post-stimulation when the signal decreases below 63% of the maximum post-stimulation value. Error bar and span: Mean +/-SEM.

Particularly, this experimental paradigm evoked a large and consistent photostimulation-driven response in the LFP signal (Figure 2B), suggesting that the photostimulation evoked a large-scale synaptic input at the neuronal region which received 4-AP. Thus, this photostimulation functions as a test of this population’s response to a strong, synchronous input. Indeed, we found that 78% of ictal onsets in these experiments were timed at 0 secs relative to photostimulation trials (Figure 2C, p(V test for non-uniformity) = 1.3e-4), suggesting that many seizures were directly induced by photostimulation. This effect may result from the optogenetic stimulus contributing to a synergistic positive feedback loop driving increased excitability of networks towards seizure onset *in vivo* (49).

The simultaneously collected LFP signal was used to classify the post 4-AP injection experimental period into ictal (classified between seizure onset and termination) and interictal (classified as any period outside of the ictal period) states. We then compared the photostimulation responses before 4-AP injection (baseline) and after 4-AP injection (interictal). Under baseline conditions, photostimulation-evoked response in the mean FOV GCaMP signal demonstrated the large-scale magnitude of widefield optogenetic stimulation (Figure 2E, F; mean stdev;/-+ response magnitude, 0.22 dFF +/-0.14; decay constant, 0.47 sec +/-0.25).

Interestingly, we found a significant increase in the photostimulation-evoked calcium response magnitude and the decay constant of the post-stimulation calcium trace in the interictal state (Figure 2E, F; mean +/-stdev.; response magnitude: 0.62 dFF +/-0.66, decay constant: 0.74 sec +/-0.40; t-test (response magnitude, baseline vs. interictal): p=5e-25, t-test (decay constants, baseline vs. interictal): p=6e-18; trials within 500ms of ictal onset and offset were excluded). The increased population GCaMP response magnitude and decay constant of the photostimulated population of neurons suggests a hyperexcitability of the neuronal population in the FOV. However, this might also be explained by increased excitatory influence of the 4-AP injected region on the experimental FOV’s neuronal population.

### Single neuronal excitability in acute epilepsy

Although focal 4-AP injection directly establishes a small region acting as the seizure focus, it is still unclear whether the acutely epileptic state induced by 4-AP injection also leads to changes in neuronal excitability outside of the 4-AP injection site. Our previous widefield optogenetic stimulation experiments suggested a large-scale hyperexcitable state after 4-AP injection. However, the wide and unspecific stimulation paradigm performed *in vivo* is not able to accurately capture changes in excitability at the single neuron scale. To test this, we used all-optical interrogation in our *in vivo* experimental preparation to directly test for changes in single neuron excitability across many individual neurons simultaneously. By applying all-optical interrogation at a site distal to the 4-AP injection site, we were able to directly study how an acute focal epilepsy model affects neuronal excitability distal to the seizure focus. The simultaneously collected LFP signal provided an electrophysiological reference from the 4-AP injection site.

We used a two-photon microscope, as previously described (39, 50), with two independently scanned optical paths for simultaneous two-photon calcium imaging and holographic, two-photon optogenetic photostimulation (Figure 3A). The photostimulation optical path included a spatial light modulator (SLM) that allowed us to create desired patterns of beamlets to target individual neurons for two-photon photostimulation. We first mapped the FOV for photostimulation responsive neurons, and then selected 30-50 photostimulation responsive neurons spread across the FOV to target for photostimulation. We maintained a consistent FOV for the duration of each experiment, which allowed us to study a fixed set of targeted-neurons across the experimental conditions. We measured the photostimulation-evoked Ca^2+^ response from targeted neurons across all repeated photostimulation trials (N = 6 animals, 41.7 +/-9.5 (mean +/-stdev.) targets per subject). Under the baseline state (prior to 4-AP injection), photostimulation effectively evoked a strong Ca^2+^-response from targeted neurons on average across all trials (mean +/-stdev.: 21.3 % dFF +/-9.3). Unlike widefield photostimulation (Figure 2), there was no appreciable response on the LFP signal evoked by two-photon photostimulation (Figure 3B), indicating that our two-photon optogenetics does not lead to the propagation of widespread downstream activity. Indeed, we did not find any significant probability of ictal onset with photostimulation timing (Figure 3E, p(V test for non-uniformity) = 0.47), suggesting that our two-photon photostimulation design does not affect seizure likelihood or lead to the induction of seizure after 4-AP injection.

**Figure 3:**
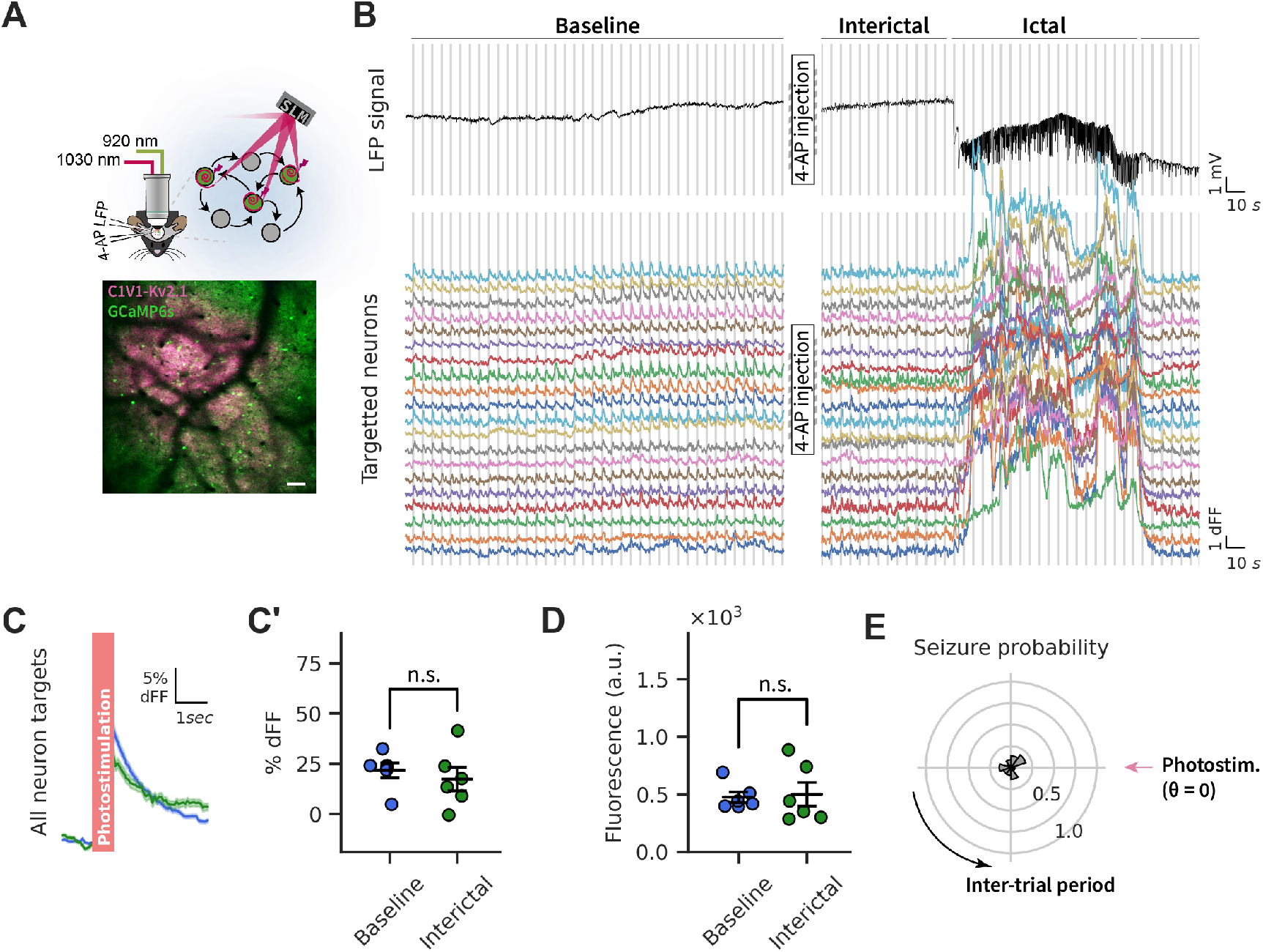
All-optical interrogation of neuronal excitability in the acutely epileptic state A. (Top) Head-fixed two-photon all-optical interrogation. (Bottom) Co-expression of GCaMP6s (green) and C1V1-Kv2.1-mScarlet (pink) at the experimental microscope FOV. Scale bar = 100 µm. B.All-optical interrogation peRF-01ormed at a fixed FOV over three states: baseline (pre-4-AP inj.), interictal (post-4-AP inj., non-seizure) and ictal (post-4-AP inj., during seizure). (Top) LFP recording from a single experiment before and after 4-AP injection. (Bottom) Calcium fluorescence traces from photostimulation targeted neurons during baseline, and the same targets during an ictal event following 4-AP injection. Photostimulation trials are represented by vertical lines over the LFP and calcium fluorescence traces. C.Photostimulation timed dFF normalised fluorescence response of photostimulation targeted neurons under baseline, interictal and ictal states (shown as mean +/-SEM across all targets in all experiments). C’: Photostimulation evoked response magnitude of photostimulation targets in each experiment (N = 6 mice, paired t-test: p=0.22) D.FOV fluorescence of photostimulation targets measured from a 500ms period before each photostimulation trial (N = 6 mice; paired t-test: p=0.85) E.The probability of seizure occurrence relative to photostimulation (binned at 1 sec intervals, N = 6 mice, p(V test for circular non-uniformity) = 0.47). SLM: Spatial Light Modulator, 4-AP: 4-aminopyridine, LFP: Local-field potential, dFF: delta F/F, mV: millivolts, n.s.: not significant. Error bars and span: Mean +/-SEM

After performing photostimulation trials under the baseline state (pre-4AP), we repeated the same photostimulation protocol in the acutely epileptic state following 4-AP injection. We measured the photostimulation evoked Ca^2+^-response from the same targeted neurons which were measured under the baseline state, allowing for a direct comparison of post-4-AP photostimulation responses to baseline measurements. This comparison showed that the average photostimulation response across all targeted neurons was not significantly altered during the interictal period (% dFF mean +/-stdev.: 17.4 +/-13.1, paired t-test (baseline vs. interictal) p=0.22). This follows the similar overall activity levels of all neurons measured in baseline and interictal periods (Figure 1F) and comparable average pre-stimulation fluorescence levels for targeted neurons (Figure 3D, paired t-test p(baseline vs. interictal)=0.85). These results suggest that in an acutely epileptic state, the single-cell excitability of neurons distal to the seizure onset zone remains at relatively baseline levels.

### Dynamics of neuronal excitability in the awake and acutely epileptic brain

The real-time activity statistics of neurons in the awake brain is a mixture of correlated and decorrelated structure (48, 51–54). The excitability of neurons is dynamically fluctuating as it integrates a balanced stream of excitatory and inhibitory inputs (55). To what extent is neuronal activity shaped by correlations in excitability? To measure this requires single-trial measurement of excitability across individual neurons. All-optical interrogation allows for a unique opportunity to measure excitability of individual cells in parallel over a large region, which reflects real-time sub-threshold dynamics by measuring the excitability of individual cells (40). The responses of photostimulated neurons are modulated by spontaneous behaviour and brain state (39).

We additionally find that the responses of individual targeted neurons show trial-to-trial variability under the baseline state (Figure 4A, mean +/-stdev.: coefficient of variation (|CV|) = 1.24 +/-0.49). Though the average excitability of neurons is not affected during the interictal state, we next asked whether the overall variability of photostimulation responses within individual neurons might be affected in the acute epileptic state. We found that the |CV| of photostimulation responses was significantly higher in the interictal state (Figure 4B, mean +/-stdev.: interictal = 2.32 +/-1.47, MWU: p(baseline vs. interictal) < 0.05, N = 6). The coefficient of variation of individual neuron responses was related to their average response magnitude and across states (Figure 4C).

**Figure 4:**
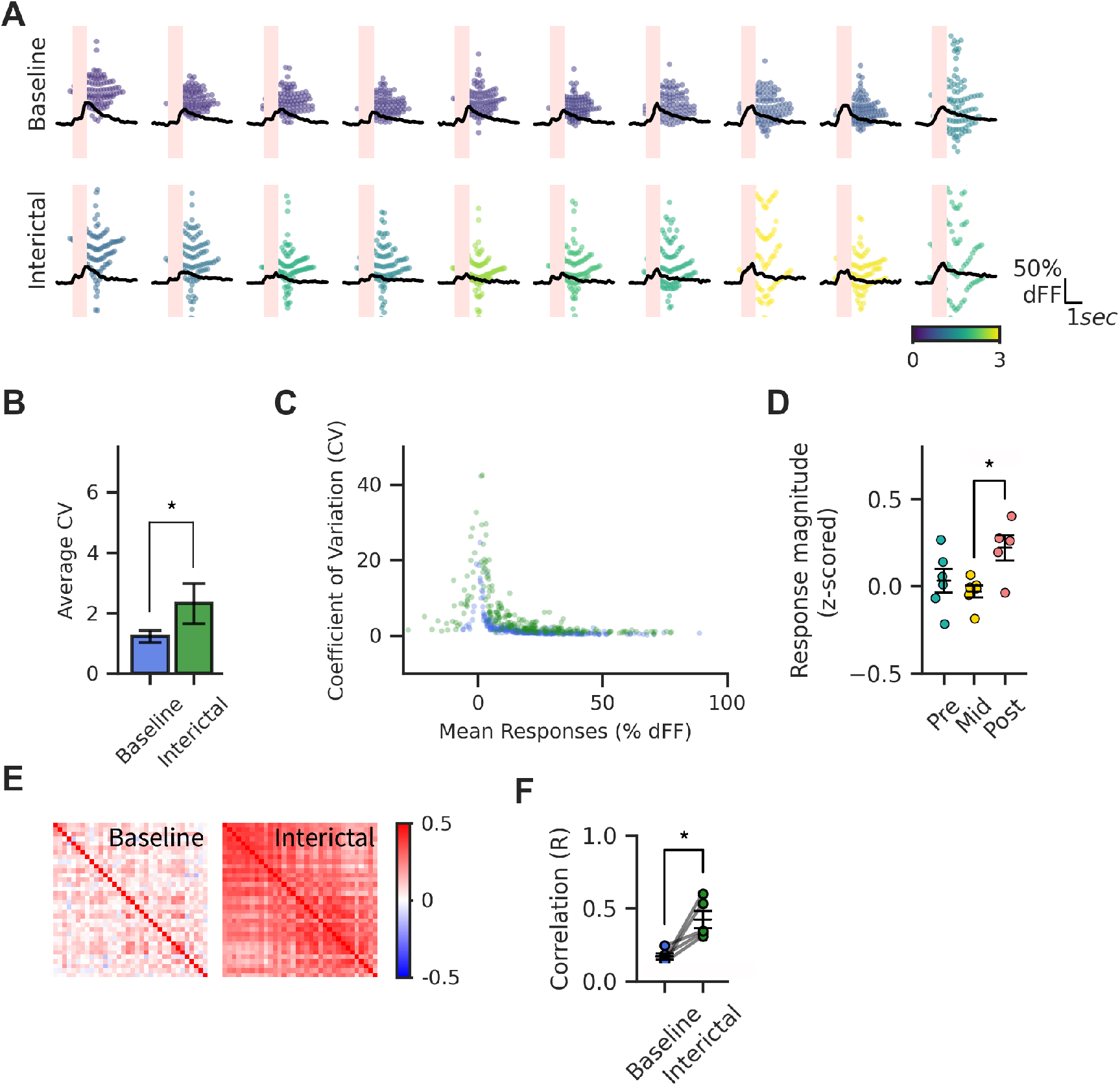
Dynamic variability of neuronal excitability across brain states A.The coefficient of variation (CV) measured trial-by-trial variability in photostimulation response magnitude of individual photostimulation targeted neurons. Representative examples of 10 randomly selected individual targets (matched Baseline (top) and Interictal (bottom) states) from a single experiment sorted by increasing CV; black trace is the mean photostimulation timed response, scatter dots are individual photostimulation responses colored by the CV of responses of each respective neuron. B.Average |coefficient of variation| (CV) of photostimulation targets under baseline and interictal conditions (N = 6 mice; *MWU: p < 0.05). C.Relationship of average photostimulation response magnitude and CV of photostimulation targets across all experiments (2-way ANOVA: p(Binned responses, 20µm bins) = 3.8e-39, p(Group:Binned responses) = 8.34e-18). Excluded datapoints: CV > 50, Mean Responses (% dFF) < -30. D.Photostimulation responses of targeted neurons (z-scored for each neuron to Baseline state) at Pre-ictal (<30secs from seizure onset), Post-ictal (<30secs after seizure termination) and middle (Interictal, outside of Pre-ictal and Postictal) (1-way ANOVA followed by a Tukey’s post-hoc test: *p < 0.05). D.Inter-neuronal cross-correlation matrix across all photostimulation targets’ z-scored responses to all trials from one representative experiment (left: Baseline, right: Interictal). Explained variance per principal component following principal component analysis dimensionality reduction of photostimulation targets’ z-scored responses to all trials (blue: Baseline, green: Interictal; 2-way ANOVA: p(Group) < 0.01, p(Group:PC) = 6.2e-6). dFF: delta F/F, CV: Absolute value of coefficient of variation. Error bars: Mean +/SEM

The trial-to-trial Ca^2+^-response magnitudes of individual neurons from baseline stimulation trials formed a normal distribution (Suppl. Figure 4), which allowed us to *z*-score the responses from each individual neuron and photostimulation trial to the distribution of responses in a target-wise manner. We also *z*-scored interictal stimulation trials in a target-wise manner to the baseline distribution of each target. This z-scoring step was essential because it normalised all response data points to a common scale, and allowed us to directly compare responses in a neuron- and trial-wise manner throughout each experiment.

In order to further investigate the source of increased variability in the interictal state, we surmised there might be a change in photostimulation responses in relation to the timing of seizures, in particular leading up to seizure onset and following seizure termination. Indeed, altered neuronal responses relative to seizure timing have been found (42), and in particular the post-ictal period displays profound, brainwide suppression on EEG (56, 57) as well as in calcium imaging (Figure 1B, Suppl. Figure 5). How is neuronal excitability affected relative to seizure onset and offset? Since, this may contribute to the variability of excitability seen in the overall interictal period, we sub-divided the interictal period into pre-ictal (0 to 30 secs leading to a seizure), post-ictal (0 to 30 secs after a seizure), and middle (interictal period outside of pre- and postictal). This sub-categorization revealed that, on average, there was no change in single-neuronal excitability relative to baseline in both the pre-ictal and middle periods. However, there was a significantly greater photostimulation response in the post-ictal phase (Figure 4D), suggesting that within the short period after seizure termination, neurons are hyper-excitable, counter-intuitive to the widespread suppression of brain activity.

The temporal (i.e. trial-to-trial) variability of single-neuron responses might be an epi-phenemenon caused by wide-scale influences on excitability affecting all neurons in our experimental FOV in parallel. To investigate this, we calculated the pairwise correlation matrix of the *z*-scored photostimulation responses of all targeted neurons across photostimulation trials from baseline and interictal (middle) states. We found there was positive inter-target correlation of photostimulation responses across trials for baseline and interictal (middle) states, suggesting that trial-to-trial variability of single-neuronal excitability within a 1mm x 1mm region is positively correlated under both states (Figure 1E). Furthermore, there was a significant increase in the correlation coefficient of target-to-target responses in the interictal (middle) state (Figure 4F, baseline: 0.17, interictal (middle): 0.42, paired t-test: p<0.05). In summary, single-neuron excitability is influenced in a moderately correlated fashion across many neurons, similar to previously described subthreshold dynamics in vivo (58) and that this can be affected in pathological states.

### Neuronal circuit excitability in the acutely epileptic state

To date, numerous all optical interrogation studies have suggested that the most prominent effect of the photostimulation of an ensemble of neurons on non-targeted neurons is an overall suppression of their activity (59–63). These studies have found a consistent, spatially-dependent effect of optogenetic photostimulation of a given neuron on the activity of surrounding non-targeted neurons within 200µm in cortex. This functional effect of photostimulated cells on neighbouring neurons is likely borne out of the reciprocal, multisynaptic anatomical connectivity of excitatory and inhibitory neurons within cortex (59, 62, 64–66). To characterise whether this structure of neural circuit excitability is affected in the epileptic state, we analysed the photostimulation-evoked responses of non-targeted neurons across baseline and acute epilepsy. We followed the approach of previous all-optical interrogation studies (39, 59, 60, 63), to measure the influence of photostimulation on non-targets in relation to their distance from the nearest targeted neuron (Figure 5A). We binned non-target to target distances into 20µm bins up to 300µm away from the target neuron.

**Figure 5:**
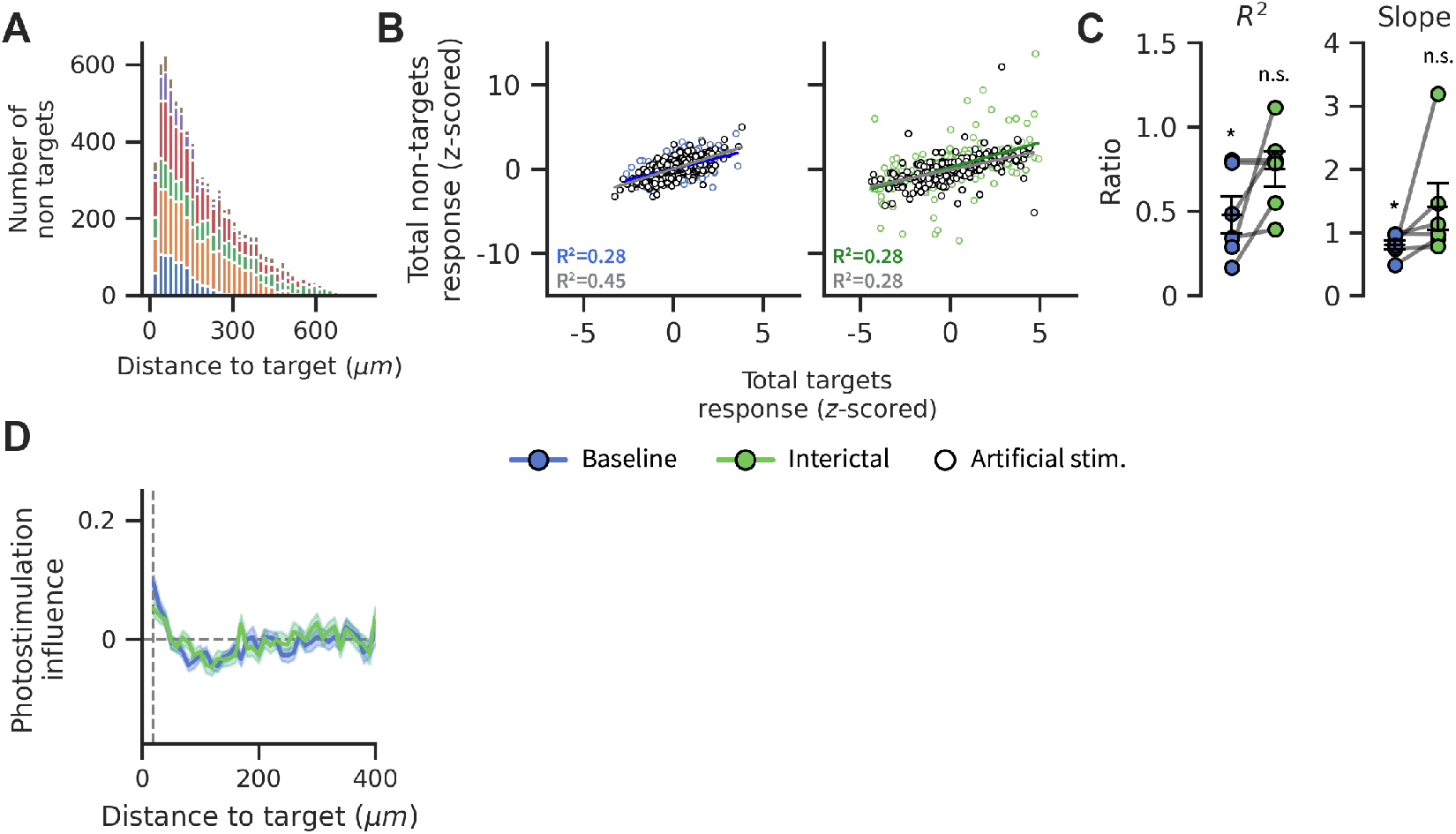
Local circuit excitability in baseline and interictal states A.Total number of measurements peRF-01ormed for distances (10µm bins) of non-targeted neurons to their nearest photostimulation target (colored by experiment, N = 6 mice). For each non-targeted neuron, the closest photostimulation target by Euclidean distance was measured. B.Relationship of single-trial total response of targeted neurons vs. total response of non-targeted neurons in baseline (left, blue) and interictal (right, green) states. Activity levels of each trial were z-scored within each experiment to allow for comparison across experiments. Artificial stimulation trials (open circles) were defined between photostimulation trials. Each point is a stimulation trial, all datapoints across all experiments are shown (N=6 mice, n = 530 baseline trials and 309 interictal trials, Pearson’s R total response targets vs. non targets: p(baseline, photostimulation) = 2.1e-39, p(baseline, artificial simulation) = 1e-70; p(interictal, photostimulation) = 7.0e-24, p(interictal, artificial simulation) = 2e-22). C.Ratio of R2 and slope of the linear regression line fitted to photostimulation trials and artificial stimulation trials for baseline (blue) and interictal (green) states (N = 6 mice, Wilcoxon signed-rank test, two-tailed comparison for different than 1, *p < 0.05). D.The influence of photostimulation on the activity of non-target neurons (n = 8567) relative to their distance (10µm bins) from the nearest target neuron under baseline (blue) and interictal (green) states (2-way ANOVA: p(Distance) = 5.5e-51, p(Group) = 0.80, and p(Group:Distance) = 0.35). n.s.: not significant. Error bars and spans: Mean +/-SEM.

In order to measure the influence of a photostimulation protocol on non-targeted neurons, it is necessary to first define an expected activity level for each neuron in the absence of photostimulation (59). The influence of photostimulation on a given neuron is defined as the difference in photostimulation-evoked response relative to its expected activity level in the absence of photostimulation. In order to define this expected activity level, we rely on the observation that the excitability of neurons within a population is strongly positively correlated. We then use this relationship to approximate the real-time expected activity level of any given individual neuron based on the overall activity state of the experimental population. We further test this relationship using the total response of a photostimulation-targeted ensemble to the total response of all non-targeted neurons in a trial-wise manner (Figure 5B). Both response measures were *z*-scored trial-wise within each experiment to allow for comparison across all experiments. There is indeed a positive relationship between the total response of a photostimulation-targeted and non-targeted neurons (Figure 5B, R²(baseline, photostimulation trials) = 0.28, p=2.1e-39), however, this could be a result of both correlated real-time excitability of all neurons or the propagation of photostimulation-evoked from targeted neurons to non-targeted neurons. To test this, we defined artificial-stimulation trials interleaved between experimentally-performed photostimulation trials (Suppl. Figure 6) and indeed found similarly that the total response magnitude of targets is positively correlated with the total response of non-targets for artificial-stimulation trials (Figure 5B, R²(baseline, artificial trials) = 0.45, =1.0e-70). In fact, the ratio of the correlation strength (R² and slope) of experimental-photostimulation trials to artificial-stimulation trials was significantly lower than 1 during baseline (Figure 5C), suggesting that photostimulation generates a negative gain on the relationship between activity of targeted and nontargeted neurons.

In comparison to the baseline state, the interictal state also maintained a strongly positive relationship between photostimulation evoked targets activity and non-targets activity, including for artificial stimulation trials (Figure 5B, R²(interictal, photostimulation trials) = 0.28, p=6.9e-24; R²(interictal, artificial trials) = 0.28, p=2.3e-22). However, notably, the ratio of correlation (R² and slope) of experimental-photostimulation trials to artificial-stimulation trials was no longer significantly different than 1 (Figure 5C). This suggests that the overall inhibitory influence of targeted neurons to non-targets is weakened in the interictal state.

After demonstrating a significant positive relationship between the activity of targeted and non-targeted activity, we may calculate the effect of photostimulation on a single trial as the photostimulation-evoked response z-normalised to the distribution of responses formed by the overall experimental population’s response on a given trial, where this is an approximation of the expected activity of all neurons for the same trial. We used this measure to calculate the influence of the photostimulation of a given target neuron on the nontargeted neurons located within a 300µm radius. We found, in line with previous reports (59–62), that the influence of photostimulation of a given neuron is positive for nearby neurons, and negative for neurons up to 200µm away (Figure 5C). Interestingly, this influence profile with respect to distance was maintained during the interictal state, and there was no significant difference in the influence measure between baseline and interictal (2-way ANOVA; distance to target: p< 1e-27; trial state: p> 0.05).

### Spatial profile of single-neuronal and local-circuit excitability during seizure propagation

Our analyses have thus far demonstrated that all-optical interrogation captures a dynamic landscape of excitability in the awake brain and during the acutely epileptic state. Next, we studied how excitability at single-neuron resolution across multiple neurons is affected during seizure propagation, which in our model, begins in a localised zone and propagates through the experimental FOV. While many previous experiments have studied single-cell excitability during seizure propagation, this has remained difficult to assess in the *in vivo* setting with singlecell resolution. Specifically, the presence of widespread inhibition in the ictal penumbra has been predicted during the process of seizure propagation (10, 11, 13–15, 67).

From calcium imaging, we first identified the location of the seizure wavefront in all stimulation trials which were classified to have occurred in the duration of a seizure (Suppl. Figure 7). Each targeted neuron was then classified based on its location inside or outside of the seizure wavefront on each photostimulation trial (see methods). Although the average photostimulation response was lowered in these neurons during the ictal period (Figure 3F), the Ca^2+^-signal during this period is difficult to interpret since seizing neurons show a raised Ca^2+^-signal due to intense, tonic activity during an ongoing seizure. The average responses of targets inside the seizure boundary are likely confounded by a potentially saturated pre-stimulation activity level or fluorescent calcium signal as a virtue of those neurons being part of the seizure territory. In contrast, the average responses of targets outside of the seizure territory were not significantly different compared to baseline or the interictal conditions (Suppl. Figure 7). The average pre-stimulation fluorescence signal was not significantly different for targets outside of the seizure boundary compared to baseline or interictal conditions, thus neurons outside of the seizure territory are not confounded by a raised activity level or fluorescent calcium signal. Therefore, for further analysis, we focused on the responses of neurons that were classified outside of the seizure wavefront boundary.

We next measured how the photostimulation responses of a given targeted neuron located outside of the seizure is related to its distance from the seizure wavefront. We again *z*-scored responses for each target across all photostimulation trials to the baseline distribution in a targets-wise manner. There was a significant relationship of the photostimulation responses of targeted-neurons in relation to their distance from the seizure wavefront (Figure 6B, 1-way ANOVA: p=1.2e-12; 64 024 total measurements, n = 154 stimulation trials with seizure wavefronts, N = 6 experiments). Specifically, photostimulation responses of targeted neurons are suppressed in close proximity to the seizure wavefront (< 100µm), while being increased distal to the seizure wavefront (> 100µm) relative to their baseline responses. We also measured the non-specific neuropil signal derived from all neuronal ROIs and found a significant relationship in relation to the seizure wavefront (Figure 6B, 1-way ANOVA: p<10e-5). Interestingly, the spatial relation of the neuropil signal to the seizure wavefront was opposite to the photostimulation responses of targeted neurons, wherein the neuropil signal of neurons decreases further away from the seizure wavefront.

**Figure 6:**
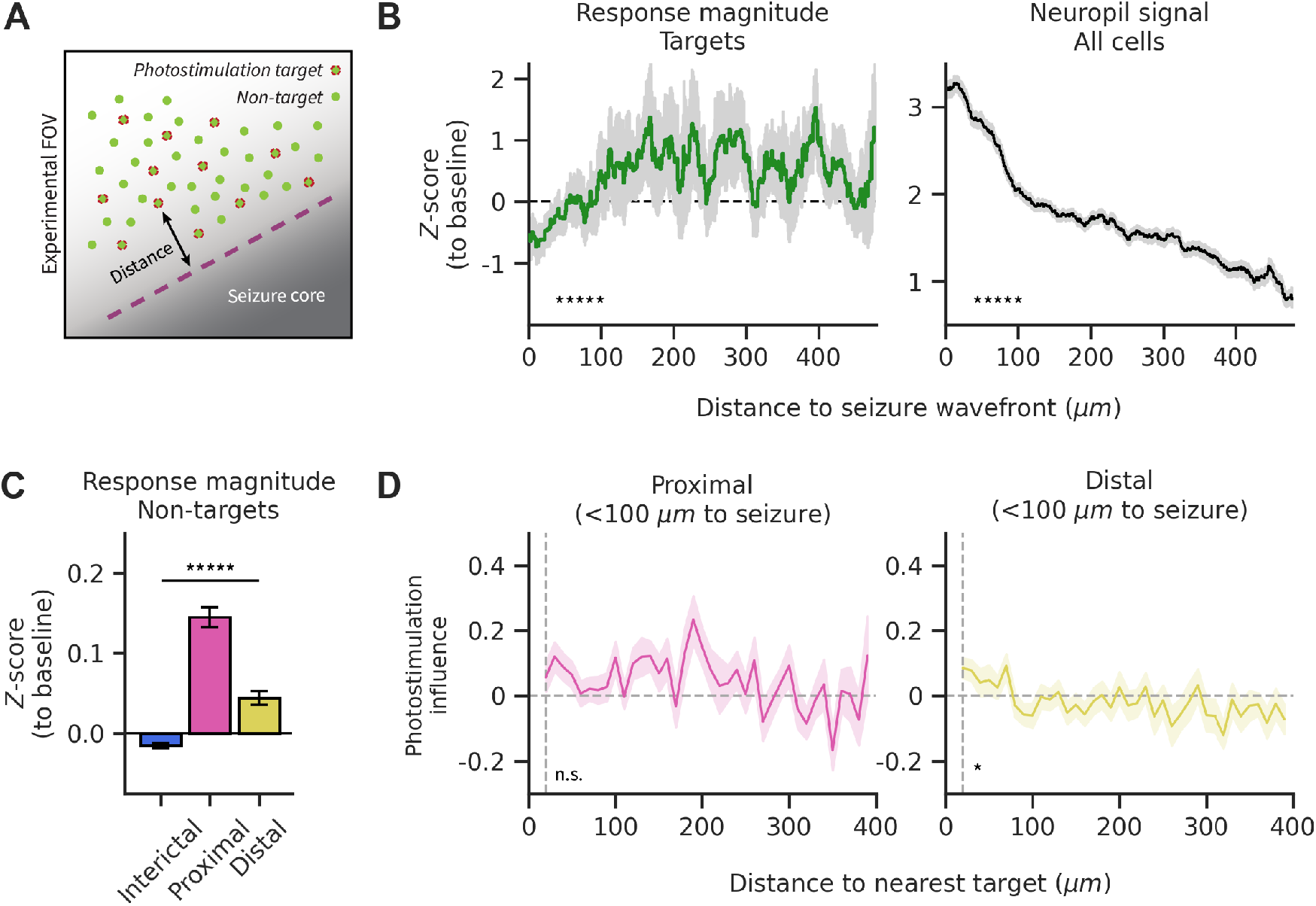
Neuronal and local-circuit excitability during seizure propagation A.Distance to seizure wavefront (dashed line) was measured linearly for photostimulation target cells (red + green filled) and non-targeted cells (green filled only) at each photostimulation trial during seizure spread inside the FOV. B. (left) Profile of photostimulation responses of targeted neurons over distance (20 µm rolling bins) relative to seizure wavefront. Photostimulation responses are z-scored to each target’s distribution of baseline responses (1-way ANOVA: *****p<1e-11). (right) The average neuropil signal associated with all cells over distance (20 µm rolling bins) relative to seizure wavefront. Neuropil measurements are taken at photostimulation trials (from 0 ms to 250 ms prior to stimulation) and z-scored to each cell’s distribution of baseline measurements (1-way ANOVA: *****p<1e-5). C.Average photostimulation response of non-targeted neurons during the interictal state (blue), and classified proximal (pink) and distal (gold) to the seizure wavefront (N = 6 mice; 1-way ANOVA: *****p < 1e-46). D. Influence of photostimulation on the activity of non-targeted neurons proximal (left) and distal (right) to the seizure wavefront (1-Way ANOVA: p(Proximal) = 0.17, *p(Distal) < 0.05). Error bars and spans: Mean +/-95% confidence interval (A, B); Mean +/-SEM (C; D).

Finally, we also measured local-circuit excitability by quantifying the responses of non-targeted cells during seizure propagation. Based on our earlier observation that the photostimulation response of neurons is related to their distance from the seizure wavefront, we divided non-targeted neurons into two groups: proximal (<100µm) to the seizure wavefront and distal (>200µm) to the seizure wavefront. We find that the mean photostimulation response of proximal neurons is significantly higher than the response of distal neurons, and both are also significantly higher than responses during the interictal state (Figure 6C, 1-way ANOVA: p=8.2e-47, proximal vs. distal: p=2.0e-10, interictal vs. proximal: p=2.8e-12, interictal vs. distal: p=2.1e-10). We subsequently measured the influence of photostimulation on non-target neurons (influence calculation as previously described) using expected activity normalisation from only non-target neurons classified outside of the seizure for a given trial. This showed that non-target neurons proximal to the seizure wavefront show no statistically significant relationship between the photostimulation influence and the distance to nearest-photostimulation target (Figure 6D, 1-way ANOVA: p=0.17). However, distal neurons maintained a statistical relationship between the photostimulation influence and distance to nearest-photostimulaion target (Figure 6D, 1-way ANOVA: p=0.02).

Overall, all-optical interrogation performed during the propagation of seizure demonstrates a profile of single-neuronal and local-circuit excitability that is significantly related to the relative location of seizure away from a given set of neurons.

## Discussion

Until now, how neuronal excitability across a neuronal population changes during seizure propagation remained largely unknown. We leveraged the use of all-optical interrogation to measure changes in neuronal excitability at the scale of individual neurons in an acutely epileptic state. Our study shows a mixed profile of hyper- and hypo- excitability of neurons based on their distance from the seizure wavefront.

The assaying of neuronal excitability across states has classically been the domain of single-cell electrophysiological studies, and indeed studies performed *in vivo* levy significant insights into the behaviour of neurons in this setting (1). There is a paucity of methods to assay neuronal excitability in the *in vivo* setting to study subthreshold dynamics that give insights into the integration of excitatory and inhibitory inputs of neurons. Valero et al., (2022) combined optogenetic micro-stimulation and multi-unit electrophysiological recording to measure the dynamic excitability of neural assemblies *in vivo* in the freely-behaving setting, allowing for the discovery of increased and decreased states of excitability of CA1 cells in place fields and during sharp-wave ripples, respectively (68). However, this method was not designed to allow for excitability measurements of individually targeted neurons. In the present study, we present a new approach to measure single-neuronal intrinsic excitability and population-scale excitability using all-optical interrogation during pathological brain activity. This allowed us to target stimulation and recording at single neuron resolution across a cortical neuronal population.

We also show here changes in wide-scale excitability across neurons over long timescale experiment durations, wherein there is substantial interneuronal correlation in trial-to-trial variability of excitability under baseline. Interestingly, this interneuronal correlation was further substantially increased after establishing the acutely epileptic state, despite the average magnitude of single-neuron stimulation responses remaining at baseline levels across all trials.

Widefield optogenetic stimulation of a large scale population showed that the acutely epileptic brain is highly sensitive to large, synchronous inputs. We found that seizure induction was strongly associated with widefield stimulation, however notably the majority of stimulation trials did not result in seizure induction. This suggests that the overall network displays a dramatic, all-or-nothing type of shift in network excitability at certain moments of time when the likelihood of seizure onset becomes heightened (49). The focal 4-AP injection epilepsy model created a localised region of epileptic hyperexcitability, and photostimulation of a distal neuronal population in the acutely epileptic state evoked activity responses of higher magnitude and longer decay constant. Since the opsin-expressing population of neurons was not exposed to 4-AP, this likely occurs from increased propagation of activity between the photostimulated-neuronal population and the hyperexcitable 4-AP-neuronal population. These population-scale findings support the suggestion that longrange networks in the brain become hyperexcitable within the epileptic state (29).

In contrast with the findings during population-scale stimulation, when the same neuronal population was stimulated with holographic photostimulation aimed at 30-50 individual neurons, there was no increase in the photostimulation-evoked response of the acutely epileptic state. Furthermore, simultaneous stimulation of tens of neurons (a scale of cells which has been shown previously to be sufficient to drive behavioural perception in the mouse brain (60, 69–71), there was no association with the induction of seizures. Thus, single-neuronal excitability is not affected in the acutely epileptic state, despite an increase in the population-scale excitability.

Instead, all-optical interrogation found a change in the response statistics of photostimulated neurons. The photostimulation responses of neurons became significantly more variable in the acutely epileptic state. We investigated the heterogeneity in the response of neurons preceding seizure onset and following seizure termination. We found that there was no significant increase in single-neuronal excitability preceding seizure onset, a timeframe in which the population activity dimensionality is proposed to be decreased (42). Surprisingly, we also found that single-neuronal excitability was increased following seizure termination. The post-ictal state is classically thought to be a hypo-active state (57, 72, 73), and indeed we also find an associated temporary suppression of widescale GCaMP signal (Suppl. Figure 5). This suggests that the increased photostimulation-evoked responses of individual neurons might be attributable to a post-synaptic low-conductance state. These data add a crucial constraint to the mysterious process of seizure termination (74, 75).

A seizure is an extreme state of neuronal activity propagation at the population scale. This is a striking departure from the physiological operation of a system such as cortex where excitation/inhibition balance normally dictates a net inhibitory effect of single-neuronal input (59). The seizure wavefront defines a striking boundary of highly active (seizing) and inactive (non-seizing) neurons that moves as the seizure recruits further neuronal populations. How does a propagating seizure influence the excitability of neurons not-yet-recruited into the seizure?

Contrary to current models of seizure, we find that neurons in the ictal penumbra show increased neuronal excitability, but that there is also a decrease in neuronal excitability in close proximity to the seizure (<100µm). Furthermore, the response of non-targeted neurons to photostimulation of nearby targeted neurons is greater for non-targeted neurons in close proximity to the seizure wavefront than for non-targeted neurons which are distal to the wavefront. If there is decreased neuronal excitability near the seizure, then why would the responses of non-targeted neurons be facilitated in the same region? A mechanism that could explain both of these findings is shunting inhibition. Shunting inhibition divides a depolarizing current input during moments of high neuronal conductance created by high levels of synaptic inputs, in spite of balanced excitatory and inhibitory neurotransmission (35, 76).

Shunting inhibition has a significant inhibitory influence on current inputs at the soma, such as our somatically-targeted 2P photostimulation method (77). Near the seizure wavefront, there is intense activation of both excitatory and inhibitory cells: the former would lead to significant activation of synapses targeting dendrites of local excitatory cells, and the latter includes peri-somatic targeting inhibitory cells (PV^+ve^ cells) which can cause a strong shunting inhibition effect (78). Therefore, whereas somatically-targeted photostimulation inputs are highly susceptible to shunting inhibition, depolarizing input at dendrites likely does not experience the same magnitude of inhibition. Furthermore, the functional effect of activating somatostatin interneurons - the dominant inhibition at dendrites - appears to be to desynchronize excitatory neuron activity, rather than to inhibit neuronal activity (51). Since shunting inhibition is directly correlated to *E*_*GABA*_ and requires intense GABAergic neurotransmission, it is likely that the depolarization of *E*_*GABA*_ during seizure propagation occurs in the post-synaptic somatic targets of the most active inhibitory neurons located within the seizure wavefront. Notably, a complete reversal of *E*_*GABA*_ in a given neuron that which leads to supra-threshold GABAergic neurotransmission (79) - does not occur until that neuron is recruited into the seizure (20), allowing for an intermediary phase of depolarized *E*_*GABA*_ where a primarily shunting effect is present (80). A transformation of dendritic excitability is found to occur at seizure initiation (49), however it’s unclear if such a transformation also occurs in neurons outside of the local region of hyperexcitability. The capability of diffraction-limited optogenetic stimulation and imaging of all-optical interrogation allows for future experiments to measure dendritic excitability to answer if there is indeed a subcellular difference in the excitability of neurons in the seizure state.

Our results suggest a refinement to the model of seizure propagation which has been indicated to date. Our results demonstrate that there is a decrease in the neuronal excitability of excitatory neurons within 100µm of the seizure wavefront. This distance corresponds with the electrophysiologically-mapped connection probability of inhibitory neurons → excitatory neurons (81, 82). This suggests that the source of inhibition for neurons with decreased excitability in seizure primarily originates from inhibitory neurons located directly at the seizure wavefront - i.e. the location of maximum tonic activity (7). This is supported by the close correlation in the rise of a given inhibitory neuron’s recruitment into seizure and the surrounding local region’s recruitment into seizure. This is distinct from previously proposed models in which inhibitory neurons located in the ictal penumbra, which is spatially distal to the seizure wavefront, are imparting inhibition onto their synaptic partners and lowering excitability in the ictal penumbra. We find instead that the excitability of neurons in the ictal penumbra is increased, whereas neurons close to the seizure wavefront show decreased excitability.

Our method of measuring excitability is not able to directly determine the mode of inhibition - hyperpolarizing vs. shunting inhibition - that causes the inhibitory effect on excitability, since this would require intracellular measurement of *E*_*GABA*_. We propose that shunting inhibition plays a significant role in this setting given that the seizure wavefront is intensely active and the locally connected post-synaptic neurons would experience a significantly raised conductance. Indeed, many studies find depolarized *E*_*GABA*_ in epilepsy and seizures (20, 32), which increases the ratio of shunting to hyperpolarizing inhibition. Whereas usually a single-neuron’s influence on its local neighbours is simultaneously both facilitatory and depressing, relative to inter-somatic distance (59), during seizure propagation, photostimulation responses are only depressed within this inter-somatic distance, further lending strength to the proposed significance of strong somatic shunting inhibition. This inhibition mode might also help explain why neuronal excitability is increased following seizure termination: we found that the widescale calcium signal was lower immediately following seizure termination, suggesting that the number of synaptic inputs in the imaging region is decreased, thereby also decreasing membrane conductance and lowering shunting inhibition.

Seizures are states of hyper-activation and hyper-excitation, therefore naturally they must involve the failure of inhibition in some form. All-together, we propose that the spatial structure of inhibition during an evolving seizure is more closely consistent with a localised pattern, rather than the previously suggested feed-forward pattern. It remains mysterious that regions outside of the local influence of neurons at the seizure wavefront show little neuronal firing during seizure, despite our findings suggesting raised neuronal excitability.

## Methods

### Animals and surgical preparation

Animal experimentation was carried out in accordance with the guidelines and regulations of the UK Home Office (Animals in Scientific Procedures Act of 1986) and the University of Oxford Animal Welfare and Ethical Review Board. Adult mice of both sexes (2-6 months of age) on a C57BL/6 background were used for experiments. Four wildtype and seven GCaMP6s transgenic mice (B6;DBA-Tg(tetO-GCaMP6s)2Niell/J x CamK2a-tTa(AI94)) were used in this study.

To prepare mice for imaging experiments, mice underwent a single surgery consisting of headplate implantation, cranial window implantation and viral injection. Mice were anaesthetized with isoflurane (5% for induction and 1%-2% for maintenance), and subsequently mounted into a heated stereotaxic frame. A perioperative injection of 0.1 mg/kg buprenorphine (Vetergesic), 5 mg/kg meloxicam (Metacam) was administered. The scalp was sterilised using chlorhexidine gluconate and isopropyl alcohol (Chlo-raPrep) and subsequently a midline incision was made to expose the skull. After removing the scalp bilaterally, the location of the cranial window implant was stereotaxically marked (right somatosensory cortex: -1.9mm AP, +3.8mm ML from Bregma). Then, a custom-machined aluminium headplate with a 7mm imaging well was secured on the skull using dental cement (C&B Metabond). A 3mm circular craniotomy was drilled centred on the previously marked location and the skull removed after soaking with saline. The dura overlying the exposed cortex was also carefully removed under regular saline flushing. After achieving control over any actively bleeding blood vessels, an 800nl injection of a diluted virus mixture was performed. The viral injection mixture consisted of C1V1-Kv2.1 (AAV2/9-CaMKIIa-C1V1-t/t-kv2.1-mScarlet; 2e12 GC/ml diluted 1:5) and—in wildtype mice—GCaMP6s (AAV2/1-syn-GCaMP6s-WPRE-SV40; 2.5e13GC/ml diluted 1:10 in sterile phosphate-buffered saline) virus was injected 300µm deep below the pial surface through a pulled glass injection needle and a hydraulic nanoinjector (Narishige) at a rate of 100nL/min; the needle was left in following each injection for 10mins to allow diffusion of virus. Finally, a cranial window, consisting of two #1 thickness coverglasses (3mm and 4mm diameter) optically adhered to one another, was implanted over the craniotomy and fully sealed using cyanoacrylate (VetBond) and Metabond dental cement. The cranial window was pre-drilled with a 800µm hole in the corner to allow access for 4-AP injection and LFP recording pipettes. This hole was offset to the edge of the cranial window to ensure the 4-AP injection and LFP recording locations were positioned away from the centre of the window where viral injection was delivered. The hole in the cranial window was filled using a silicone Kwik-cast plug before implanting over the craniotomy. Mice received buprenorphine (double check) at the time of surgery and in food gels for 2 days postsurgery. All animals were given 3 weeks to recover and reach optimal viral expression before experimentation.

### Focal 4-AP acute seizures and local field potential recording

Focal injection of 4-aminopyridine (4-AP) was used to produce repeated acute seizures of focal-onset in mice and nearby local field potential (LFP) signal was used to electrophysiologically record seizures for the duration of the experiment.

The Kwik-cast plug sealing the pre-drilled access hole of the cranial window was removed after setting up mice in headfixed setup. 4-AP (5mM) was loaded into a glass pulled pipette mounted on a micromanipulator which was then lowered into the cortex through the access hole in the cranial window under 5x widefield imaging. A second glass pulled pipette, backfilled with 45µm filtered cortex buffer solution (contain in mM: 125 NaCl, 5 KCl, 10 Glucose, 10 HEPES, 2 CaCl_2_, 2 MgSO_4_), was inserted in parallel to the 4-AP injection pipette using a second micromanipulator to approximately the same depth and location. LFP signal was collected from this pipette through a headstage, Axon 700B amplifier, a 50/60Hz noise-cancellation humbug, and digitised using PackIO. After successful insertion of both pipettes, they were not moved for the entire duration of the experiment and the Nikon 16x objective was lowered into position under water immersion to begin two-photon imaging experiments.

### Awake head-fixed two-photon calcium imaging

Twophoton imaging was performed using a resonant scanning microscope (2pPlus, Bruker Corporation), a tunable femtosecond-pulsed, dispersion-compensated laser beam (Vison-S, Coherent) and a 16x/0.8-NA water-immersion objective lens (Nikon). Total power was modulated with a Pockels cell (Conoptics) and maintained at or below 50mW on sample for all experiments. GCaMP6s and GCaMP7f imaging was performed using a 920nm beam and a GFP collection filter set, and mCherry or tdTomato imaging was performed using a 765nm beam and a RFP collection filter set. All animals were headfixed under the experimental imaging setup to confirm expression of virally-transfected calcium indicator and/or opsin at least once prior to experimentation. The cortical location for two-photon imaging and all-optical experimentation under the cranial window was selected at least 1mm away from insertion of 4-AP injection and LFP recording pipettes. Two-photon imaging was performed in Layer 2/3 at a depth of 150 to 300µm below the pia depending on optimal GCaMP6s and mCherry (for C1V1-Kv2.1) expression. For two-photon calcium imaging of interneurons in Nkx2.1-Cre-tdTomato mice, an FOV was selected at the start of the experiment at least 1mm away from the focal 4-AP injection and LFP recording site and with optimal co-expression of GCaMP7f virus with mCherry labelled interneurons. Live two-photon imaging was performed at either 15Hz (1024 × 1024 pixels; N = 4 experiments) or 30Hz (512 × 512 pixels; N = 2 experiments) and at between 0.8x to 1.5x zoom. Imaging was controlled through PrairieView (Bruker Corporation).

All experiments were terminated either when the experimental animal entered status epilepticus or the total experiment duration reached 3hrs, whichever occurred first.

### Simultaneous 1-Photon widefield optogenetics and two-photon imaging

Widefield optogenetics was performed in conjunction with two-photon calcium imaging (as described above) using the widefield excitation module of the Bruker 2pPlus. Blue light photostimulation (470nm, 10 × 10ms pulses @ 25Hz, at 3.5mW power on sample), was performed during two-photon imaging. To avoid damage to the two-photon imaging photomultiplier tubes (PMTs), highspeed shutters on all PMTs were triggered during blue light photostimulation. Triggers for blue light stimulation (sent to a Thorlabs LEDD1B LED driver) and the high-speed PMT shutters were provided to a using PackIO.

### Simultaneous two-photon optogenetic stimulation and calcium imaging (all-optical interrogation)

Two-photon all-optical experiments were performed by simultaneous two-photon stimulation of selected neurons using a spatial light modulator (SLM) and calcium imaging (as described above) in the same FOV using two independent beam paths setup in the Bruker 2pPlus. Two-photon optogenetic stimulation was performed using a pulsed fixed-wavelength fibre laser 1035nm (Monaco, Coherent) at a repetition rate of 2MHz. Multiple individual neurons were simultaneously targeted for stimulation by splitting the laser beam into beamlets using a reflective spatial light modulator (SLM) (7.68 × 7.68 mm active area, 512 × 512 pixels, Boulder Nonlinear Systems). The active area of the SLM was overfilled and the polarisation optimised for maximal first order diffraction efficiency using a half-wave plate. The zero order diffraction beam was blocked using a well bored into an optical flat using a dental drill (NSK UK Ltd).

SLM phase masks were loaded using the Blink SDK (Medowlark Optics). Phase masks were computed by applying the Gerchberg-Saxton algorithm to the xy coordinates of the targets (83). A weighted form of this algorithm was used to ensure uniform power distribution (6mW/target) across all targets to account for reduction in the first order diffraction efficiency of the SLM with increasing distance from the zero order location. An image of the SLM was relayed onto a pair of galvanometer mirrors (independent of the imaging beam path) integrated in the two-photon imaging system. The galvanometer mirrors were programmed to generate spirals of 10µm diameter by moving all beamlets simultaneously.

The affine transformation required to map coordinates from SLM space to imaging space was computed through a custom-modified version of open-source software written in MATLAB (github.com/llerussell/Naparm) and calibrated at the start of each experiment by burning arbitrary patterns into a fluorescent plastic slide. Phase masks and galvanometer voltages required to perform photostimulation were generated using NAPARM (50). Voltages were applied to the photostimulation-path galvanometers using PrairieView (Bruker Corporation). A USB data acquisition card (National Instruments) running PackIO, was used as a master synchroniser to record individual signals representing the LFP electrophysiological signal, the frame clock of two-photon imaging and various photostimulation signals (e.g. galvanometer voltages, SLM phase mask trigger and voltages).

In some experiments, a single widefield blue light stimulation trial driven using PackIO was used to test the optogenetic photostimulation-induced Ca^2+^ response while exploring the FOV and selecting the optimal region of fluorescence responsive neurons. A photostimulation-cell mapping protocol was run in all-optical experiments and online analysis of two-photon photostimulation was carried out using STAMovieMaker (https://github.com/llerussell/STAMovieMaker) to test for responsive targets. These images were used to manually select the coordinates of cells within the FOV for targeting of two-photon optogenetic stimulation.

### Analysis of two-photon calcium imaging and photostimulation experiments

Imaging data analysis was carried out in Python 3.8. Overall data analysis and procedures extended upon the framework provided in Imaging+ (github.com/Packer-Lab/imagingplus), a recently developed Python package for analysis of multimodal, time-synced, two-photon live imaging neuroscience experiments. We excluded experiments where onset/offset timings of seizure events could not be determined from the LFP recording. We also excluded one experiment where seizure propagation did not propagate throughout a significant portion of the imaging field of view.

#### Data processing

All calcium imaging movies were first processed through the Suite2p pipeline for movement correction and cell segmentation, followed by manual curation of Suite2p generated cell masks. Synchronised timeseries data signals collected as individual channels in PackIO were processed using the paq processing submodule of Imaging+. The photostimulation trigger timestamp and the total duration of the photostimulation protocol were used to define each photostimulation trial’s onset and offset. The photostimulation onset/offset timestamps were cross-referenced to the imaging frame clock signal to define photostimulation onset and offset in the imaging data. We excluded imaging frames between onset and offset of each photostimulation trial and all imaging frames from imaging trials after 4-AP injection from cell segmentation as photostimulation laser induced imaging artefacts and aberrant neuronal activity, respectively, confounded Suite2p cell segmentation performance. Fluorescent traces were recovered from photostimulated targeted neurons using a 10µm-diameter circular mask centred on each SLM target coordinate within the FOV. deltaF/F normalisation was performed on all traces.

#### Marking seizure onset and offset timing using LFP

LFP data was imported using the paq processing submodule of Imaging+. For data collected after 4-AP injection, the onset/offset of seizure was manually marked on each individual LFP recording. Seizure onset was defined as the last sharp deflection in the LFP signal before a sustained barrage of ictal discharges. Seizure offset was defined at the end of final large deflection in the LFP signal following a period of ictal discharges. All photostimulation trials outside of seizure onset/offset timings were classified as ‘interictal’, and those within as ‘ictal’. In some cases, the photostimulation experiment was initiated during an ongoing seizure, and therefore all photostimulation trials until the offset of the initial seizure were classified as ‘ictal’. No seizure onset was defined for these seizures.

#### Ictal event-photostimulation correlation

The correlation of photostimulation and ictal onset in widefield photostimulation and holographic photostimulation experiments was calculated using the pycircstat package (https://github.com/circstat/pycircstat). The interphotostimulation time period was divided into phase space of 1sec bins (0sec bin: -500ms to +500ms relative to photostimulation). The number of seizure event onset in each photostimulation phase bin was collected and the V-test for circular non-uniformity was performed with a mean direction of 0rads.

#### Measuring the seizure wavefront

The location of the seizure wavefront was manually assessed for all ‘ictal’ photostimulation trials based on a 100 frame averaged image centred on each individual photostimulation trial. Using ImageJ, two coordinates were drawn across the image to denote the boundary of the seizure wavefront. Then, the shortest distance was calculated from all neurons to the seizure boundary at each photostimulation trial.

Within each individual seizure event, the time of seizure invasion was individually marked for each photostimulation targeted neuron. This was marked at the initiation of the greatest upward deflection in the raw fluorescence signal that led to the maximum fluorescence signal during that event for that neuron. Then, a delay to seizure invasion was calculated at each photostimulation trial using this marked timing for each neuron.

#### Photostimulation responses of targeted neurons

Responses of photostimulation-target neurons on photostimulation trials were collected from circular areas defined within the FOV using the spatial photostimulation target area outputs from NAPARM. These areas were applied to the motion corrected movie output from Suite2p processing to collect the average raw calcium fluorescence signal from pixels within each target. Photostimulation response magnitudes for each target were calculated from dF/F normalised values by subtracting the average pre-photostimulation signal (from -500ms to photostimulation onset) from the mean post-photostimulation signal (from photostimulation offset to +500ms). This procedure was applied equally across all baseline and post-4AP photostimulation trials.

The variability of photostimulation responses for each neuron was calculated as the coefficient of variation of response magnitudes from all photostimulation trials belonging to the appropriate state. To generate normalised photostimulation responses for each neuron and photostimulation trial (across both baseline and post-4AP states), each photostimulation trial response magnitude was *z*-score normalised to each individual neuron’s baseline response distribution. A neuron’s baseline response distribution was derived as the mean and standard deviation of all photostimulation responses from the baseline state. With this procedure, photostimulation responses from the post-4AP state were also *z*-scored to the baseline response distribution.

To measure trial-to-trial correlated variability of photostimulation responses, a cross-correlation matrix and the average neuron-to-neuron Pearson’s correlation score was calculated from a matrix of target-neurons x *z*-score responses for each animal replicate split into baseline and interictal photostimulation trials.

#### Influence of photostimulation on non-targeted neurons

Non-targeted neurons were selected from the overall curated set of ROIs generated by Suite2p. A circular exclusion zone with radius of 20 µm was defined around the centre of each targeted neuron’s coordinate in the FOV to exclude any potential Suite2p ROIs that may have received off-target activation during photostimulation. Photostimulation responses for non-target neurons were calculated similarly to the procedure used for target neurons. To measure the relationship between photostimulation response magnitude of targets and the photostimulation response magnitude of non-targets, the total photostimulation response magnitude of all non-targets was plotted against the total photostimulation response magnitude of targeted neurons for each trial. These values were *z*-scored within each experimental replicate to allow comparisons across all replicates. A similar procedure was also performed on “artificial stimulation” trials. These were artificially created stimulation trials defined during analysis as an alternate to sham (no-photostimulation) trials. These trials were interleaved between photostimulation trials. The responses to artificially defined stimulation trials were calculated and analysed similarly to photostimulation trials to measure the relationship between normalised total response of targets vs. total response of non-targets during artificial stimulation trials.

To measure the influence of photostimulation of targeted neurons on surrounding non-targeted neurons, we used a modified version of the influence-mapping approach described by Chettih and Harvey (2019). In this method, the influence of photostimulation of targeted neurons on the activity of nontargeted neurons is quantified. In order to measure the influence imparted by photostimulation, it is necessary to first quantify a predicted level of activity for each non-targeted neuron. Then, the measured photostimulation trial response is normalized against this predicted response level to generate a photostimulation-driven influence metric for a given nontarget neuron. Based on our finding (Figure 6) that the photostimulation responses of non-targeted neurons are correlated to each other within each photostimulation trial, we used the mean photostimulation response of all non-targeted neurons as the predicted response magnitude for all non-target neurons in a trial-wise manner.

#### One-photon widefield photostimulation

1P widefield photostimulation timings were retrieved from the high-speed shutter loopback signal collected as a temporally synchronised paq channel. Although the high-speed shutters were deployed slightly before and released slightly after the LED photostimulation, there is no useful imaging data during the activation of the high-speed shutter. The coarse fluorescence calcium imaging signal was collected as the mean of the field of view from raw imaging movies. Photostimulation trial response magnitudes were quantified as the mean post-stimulation signal (collected from 0ms to +500ms post-high-speed shutter release) minus the mean pre-stimulation signal (collected from -500ms to 0ms pre-high-speed shutter deployment). The decay constant of the evoked photostimulation response was calculated as the time postphototstimulation when the mean FOV fluorescence signal decreased to below 63% of the maximum post-stimulation value. Photostimulation trials in which post-stimulation fluorescence trace did not return to below this threshold (e.g. when photostimulation evoked ictal events) were excluded from this quantification.

#### Interneuron calcium imaging

Interneurons and excitatory neurons were defined as tdTomato^+ve^ and tdTomato^-ve^ cells in virally-transfected GCaMP7f imaging in Nkx2.1-Cre::tdTomato transgenic mice. baseline imaging trials (spontaneous activity, before 4-AP injection) were processed through Suite2p to generate motion-corrected imaging data and segmented ROIs for analysis. The tdTomato image of the FOV was co-registered (using pystackreg) to the motioncorrected Suite2p output image with a transform matrix that was calculated by registering a simultaneously collected GCaMP image to the registered image output obtained from Suite2p.

While the Suite2p segmented ROIs were used to define excitatory neurons in the FOV, we used Cellpose (84) to generate inhibitory neuron ROIs from the co-registered tdTomato image. Suite2p ROIs with greater than 5% overlap in pixels with any inhibitory neuron ROIs were excluded from further analysis. Neuropil subtraction was performed on each respective Suite2p ROI’s signal. To analyse the relationship for seizure recruitment between an inhibitory neuron and its local surrounding region, an annulus with an inner radius of 20µm and outer radius of 100µm was created from the centre of the inhibitory neuron. This area was used to collect the average local signal from motion-corrected Suite2p imaging frames. Seizure recruitment for both inhibitory neuron and the local surrounding region was defined as time post-seizure onset (defined using the simultaneously collected LFP signal) to reach 65% of the maximum fluorescence signal during the seizure. To account for variability in propagation speeds across individual seizures and experiments, the recruitment delay for each neuron and its respective local signal was normalised to the mean recruitment delay of all individual local signal values.

## Author Contributions

P.T.S, T.A.V and A.M.P designed the study. P.T.S and A.M.P conducted experiments. P.T.S performed analysis with advice and supervision from T.A.V and A.M.P. P.T.S and A.M.P wrote the paper with inputs from T.A.V.

## ACKNOWLEDGEMENTS

We thank Richard Burman, Colin Akerman, Thijs van der Plas, Robert Lees, and James Rowland for useful discussion and technical support. This work was supported by funding from Wellcome Trust (204651/Z/16/Z) to A.M.P, from Vanier - Canada Graduate Scholarship to P.T.S, from Mitacs to P.T.S, from University of Toronto MD/PhD Program to P.T.S.

---

**Suppl figure 1:**
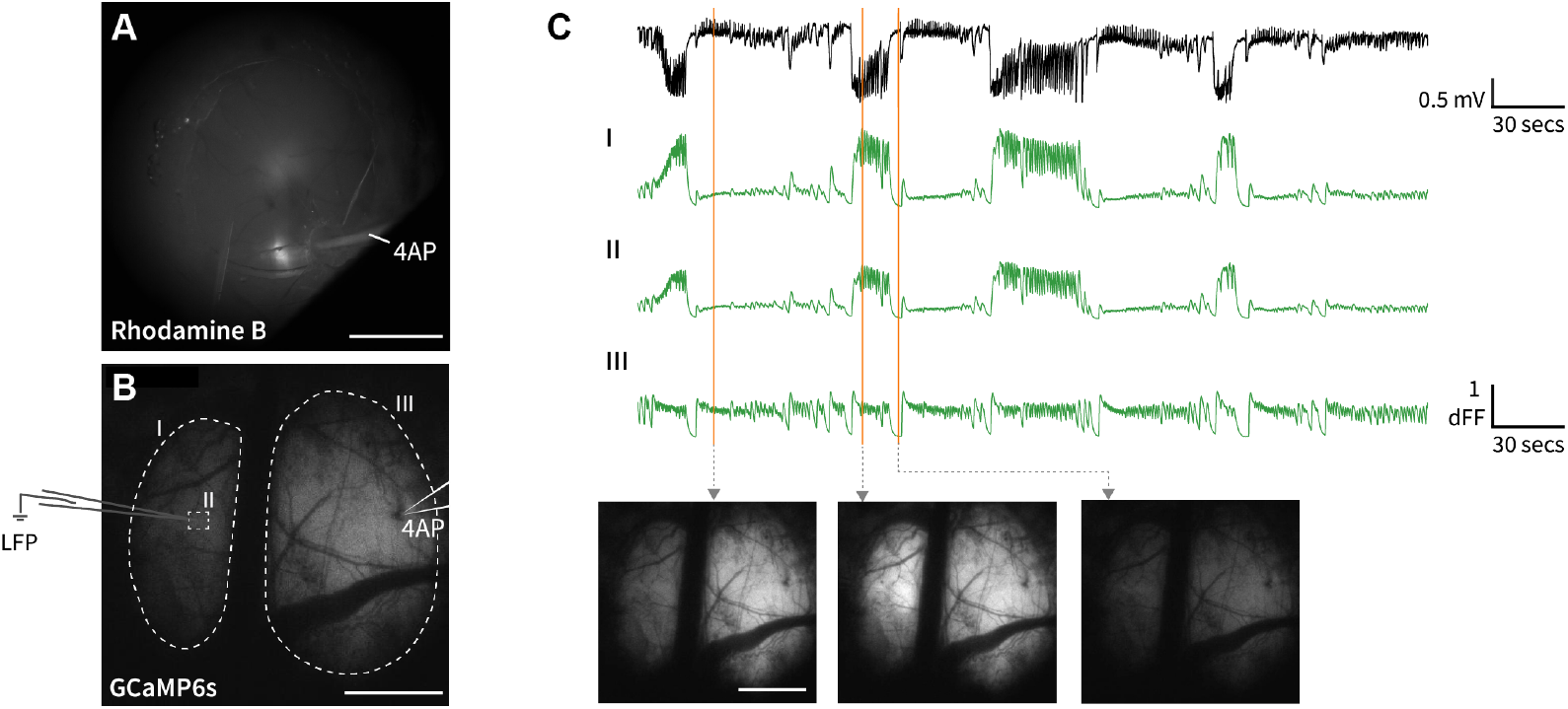
Cross-hemispheric, widefield calcium imaging of focal onset seizure A.Focal 4-AP injection (mixed with Rhodamine-B) near the edge of the implanted cranial window. Note the fluorescence signal at the centre of the cranial window in this imaging channel is from viral-injection-transduced expression of C1V1. B.Widefield, cross-hemispheric calcium imaging under a craniotomy over the frontal cortex. LFP signal was collected contralateral to the focal 4-AP injection. ROIs were selected for fluorescence signal collection relative to the 4-AP injection and LFP recording electrode (I, II and III). C.Average fluorescence signal (green traces) from various ROIs marked in panel B, synchronised to the simultaneously collected LFP trace (black trace). Below is shown single-frame widefield calcium images during interictal, ictal and following seizure termination.

**Suppl figure 2:**
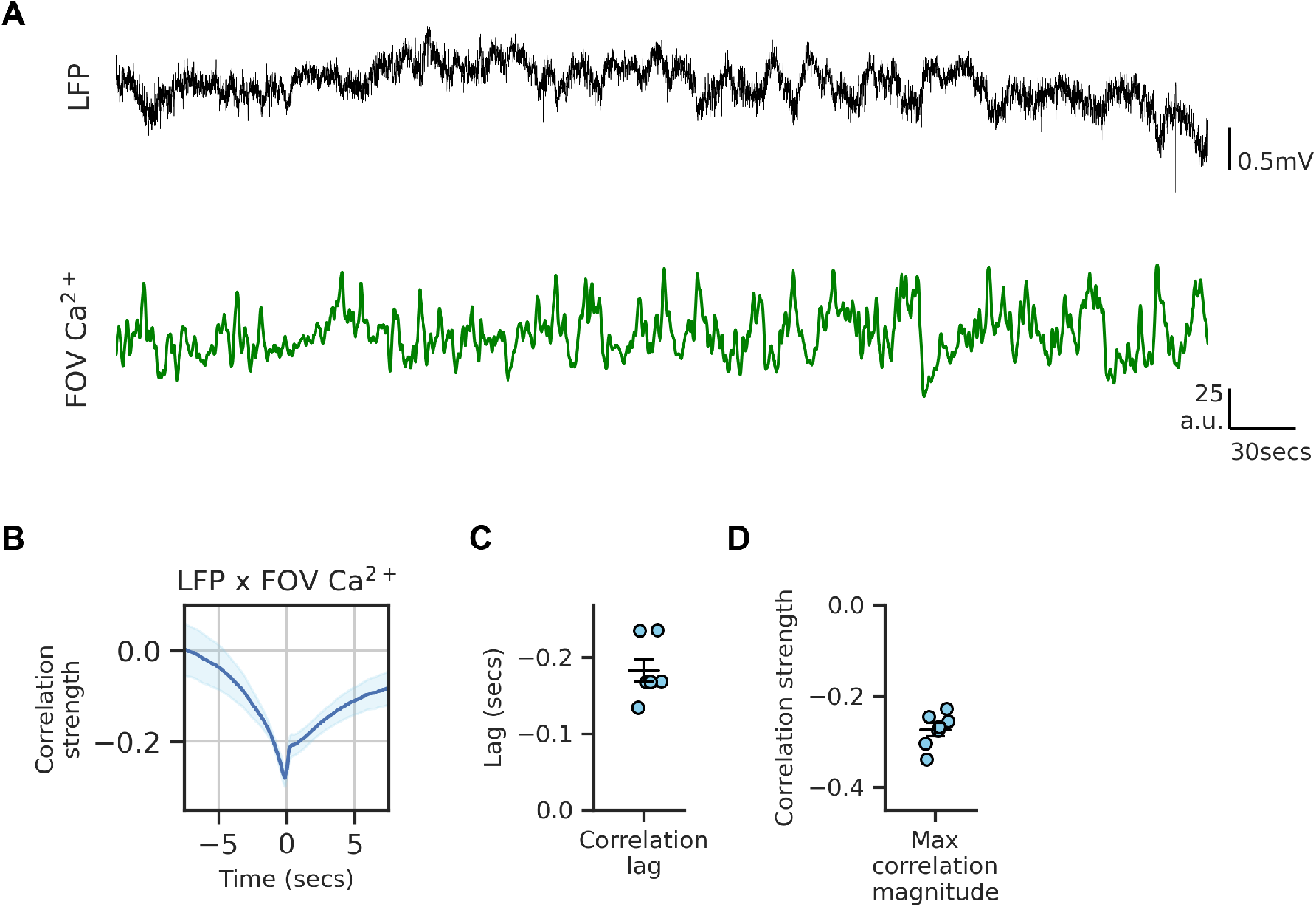
Relationship of LFP to coarse GCaMP6s signal A.Simultaneously collected LFP (top, black) and two-photon mean FOV GCaMP6s signal (bottom, green) at the same location in awake, head-fixed mice under baseline state (N = 4 mice, 15 to 60 mins of imaging per mouse). B.Cross-correlation analysis of LFP to two-photon mean FOV GCaMP6s signal. C.Lag and strength (C’) of maximum cross-correlation taken for LFP signal relative to the mean FOV GCaMP6s signal. LFP: local-field potential; Error bars and spans: Mean +/-SEM.

**Suppl figure 3:**
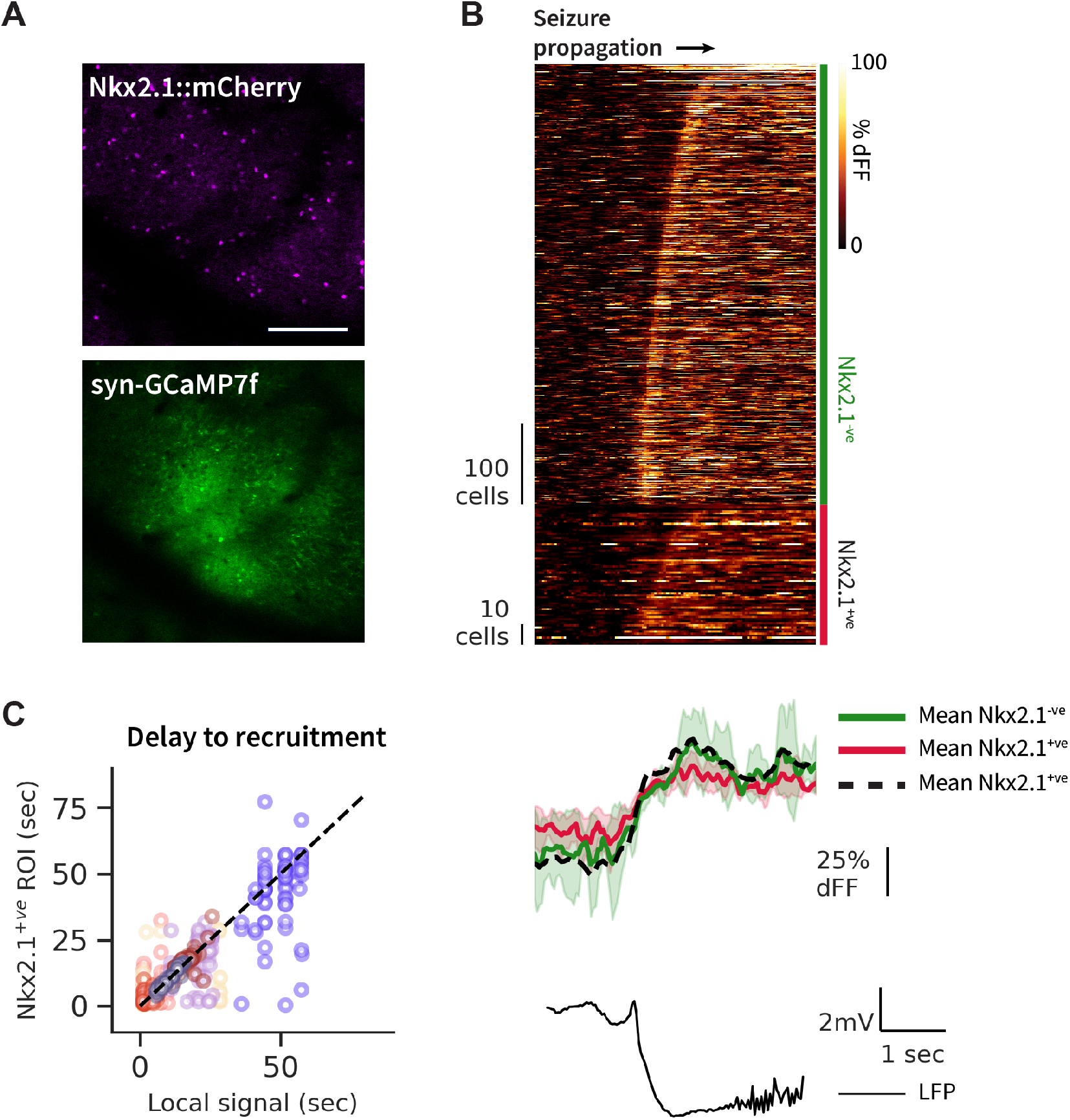
Calcium imaging of Nkx2.1-labelled inhibitory neurons at seizure onset A.Expression of virally transfected GCaMP7f (top, 920nm excitation/green emission multi-frame average) in Nkx2.1-Cre-mCherry transgenic mice (mCherry expression shown bottom, 780nm excitation/red emission multi-frame average). B.Cell-type specific GCaMP7f fluorescence signal at -1.5 to +2.5 secs around seizure onset in a representative seizure event. (left) Fluorescence signal of Nkx2.1^-ve^ (top, green bar) and Nkx2.1^+ve^ (bottom, red bar) neurons. Cells are ordered based on delay to recruitment into seizure (measured as reaching 75% of their maximal signal). Signal is normalised to the 1.5 to 0.5 secs period prior to seizure onset. (bottom) Average fluorescence signal of all Nkx2.1^-ve^ ROIs (green), all Nkx2.1^+ve^ ROIs (red), and average FOV Ca^2+^ signal (dashed black) and the simultaneously collected LFP signal (thin black) from the same timeframe around seizure onset. C.Time lag in seizure recruitment (calculated as in B) of Nkx2.1^+ve^ ROI compared to surrounding Nkx2.1^-ve^ ROIs, in relation to inter-somatic distance. Negative recruitment lag = Nkx2.1^+ve^ ROI recruited earlier than Nkx2.1^-ve^ ROI (n = 10 seizures from N = 3 mice; Kruskal-Wallis: p = 0.73). D.Cross-correlation lag and maximum correlation strength of the average signal at -2 to +2 secs around seizure onset from the FOV Ca^2+^ to Nkx2.1^-ve^ cells to (green), and for Nkx2.1^+ve^ cells to Nkx2.1^-ve^ cells (pink) (n = 10 seizures from N = 3 mice). Scale bar: 100 µm (A). Corr: Cross-correlation, FOV: Field-of-view, LFP: Local field potential, ROI: Region of interest. Error bars and spans: Mean +/-SEM.

**Suppl figure 4:**
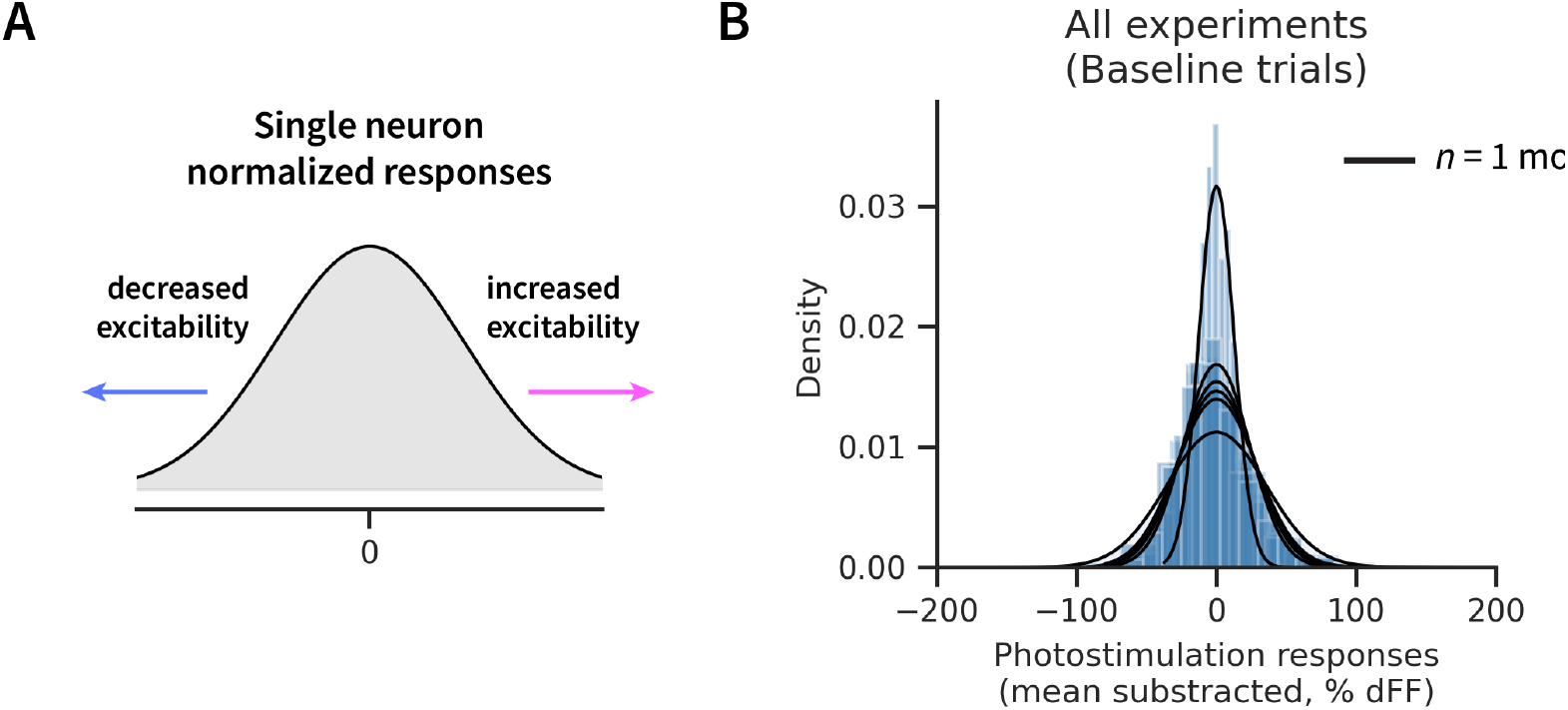
Distribution of photostimulation responses of Baseline state A.Z-score normalisation of individual neuron photostimulation trial responses allows for indexing changes in excitability of the same neuron. B.Gaussian-fitted distribution of Baseline photostimulation responses (mean subtracted) across all targeted neurons and photostimulation trials for each experiment (black). All experiments pass test for normality (***** p(for each exp.) < 1e-12).

**Suppl figure 5:**
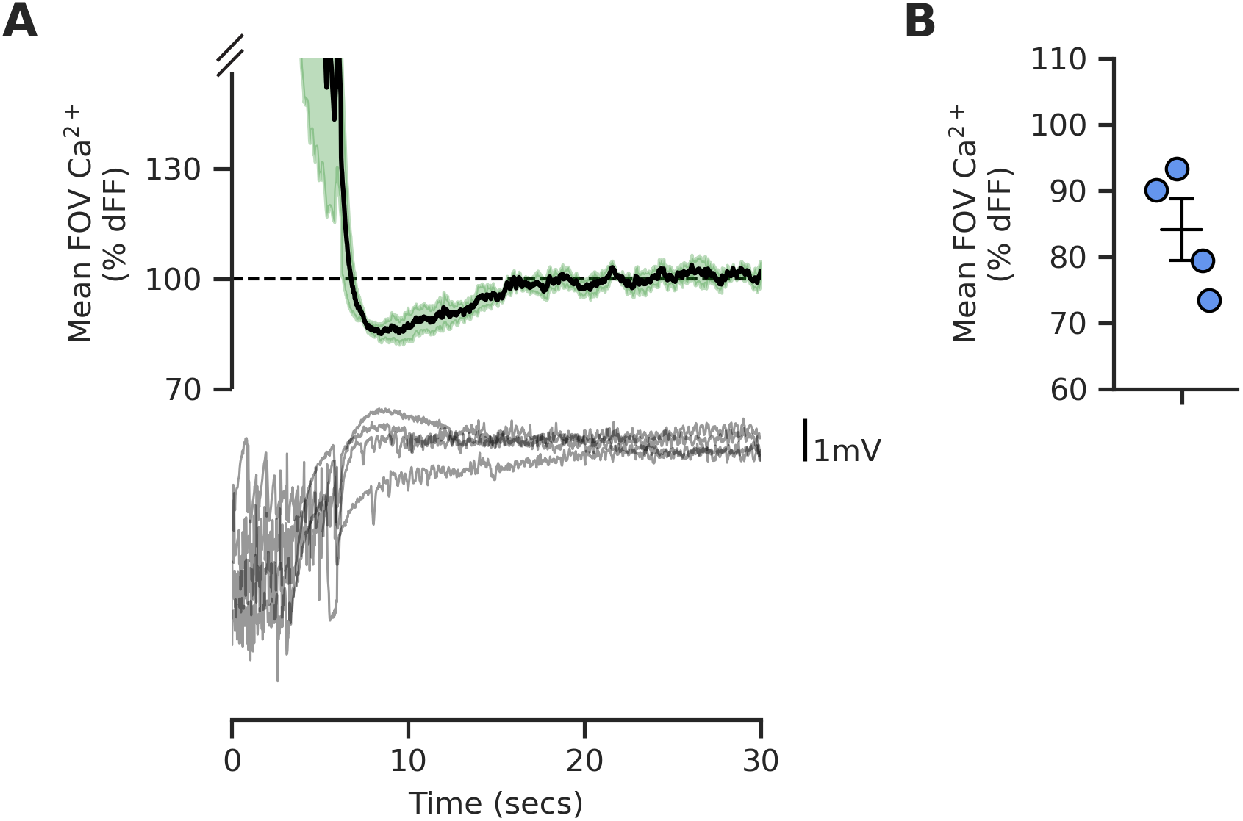
Ca^2+^ signal after seizure termination A.Dynamics of the raw FOV Ca^2+^ signal in the post-seizure termination period from a representative experiment (mean +/-SEM; normalised to the interictal period of same experiment). B.Mean (+/-SEM) of post-seizure termination normalised Ca^2+^ signal across experiments.

**Suppl figure 6:**
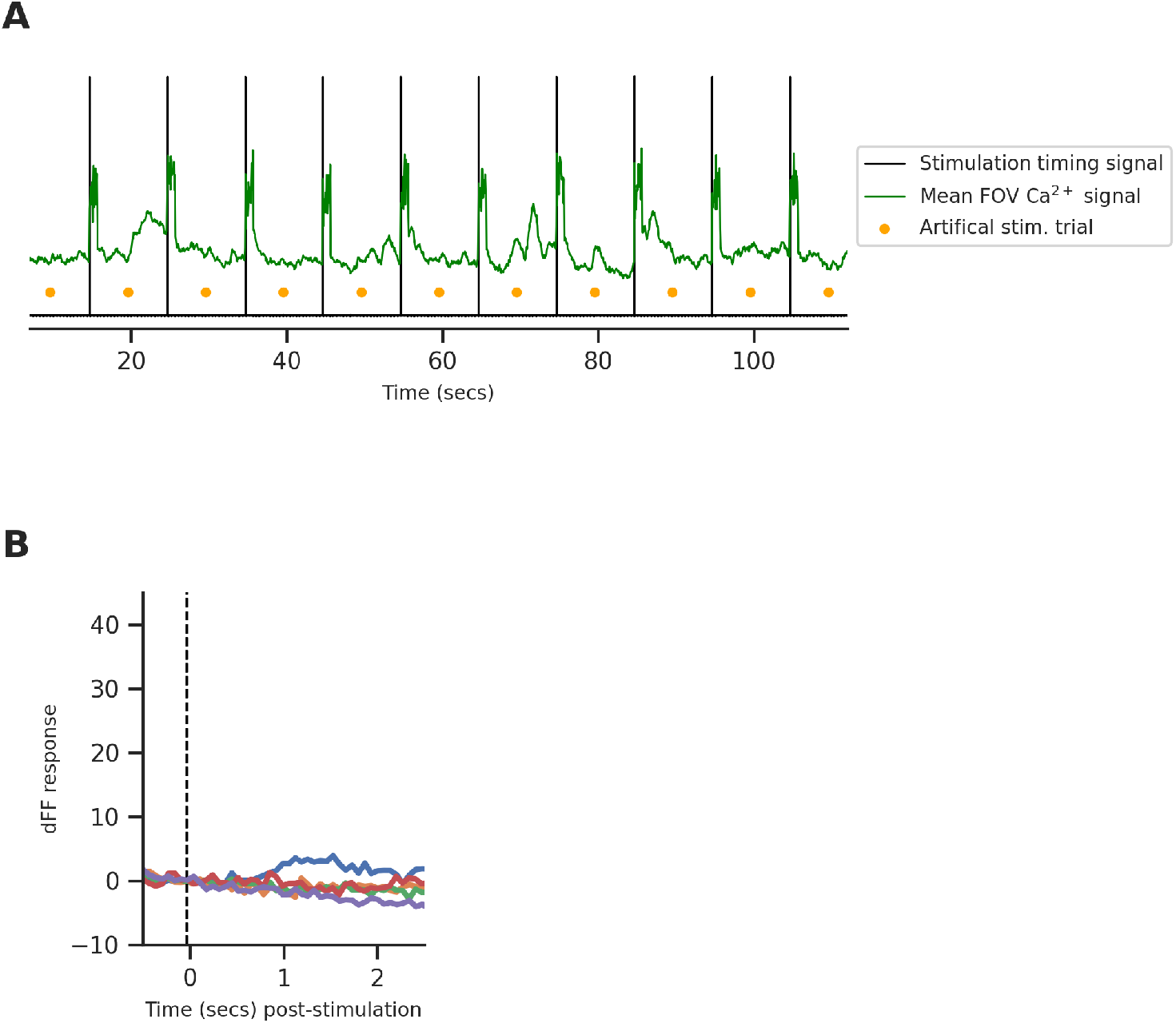
Average response of artificially defined stimulation trials A.Artificial stimulation trials were defined in between experimental-photostimulation trials for each experiment. The mean FOV Ca^2+^ signal (green) shows the imaging channel artefact generated by the photostimulation laser synchronised to the experimental-photostimulation timing signal (black), and the absence of such artefact at artificial stimulation trial times (orange markers) in one representative experiment. B.The artificial stimulation timed dFF response across all experiments from the Baseline state.

**Suppl figure 7:**
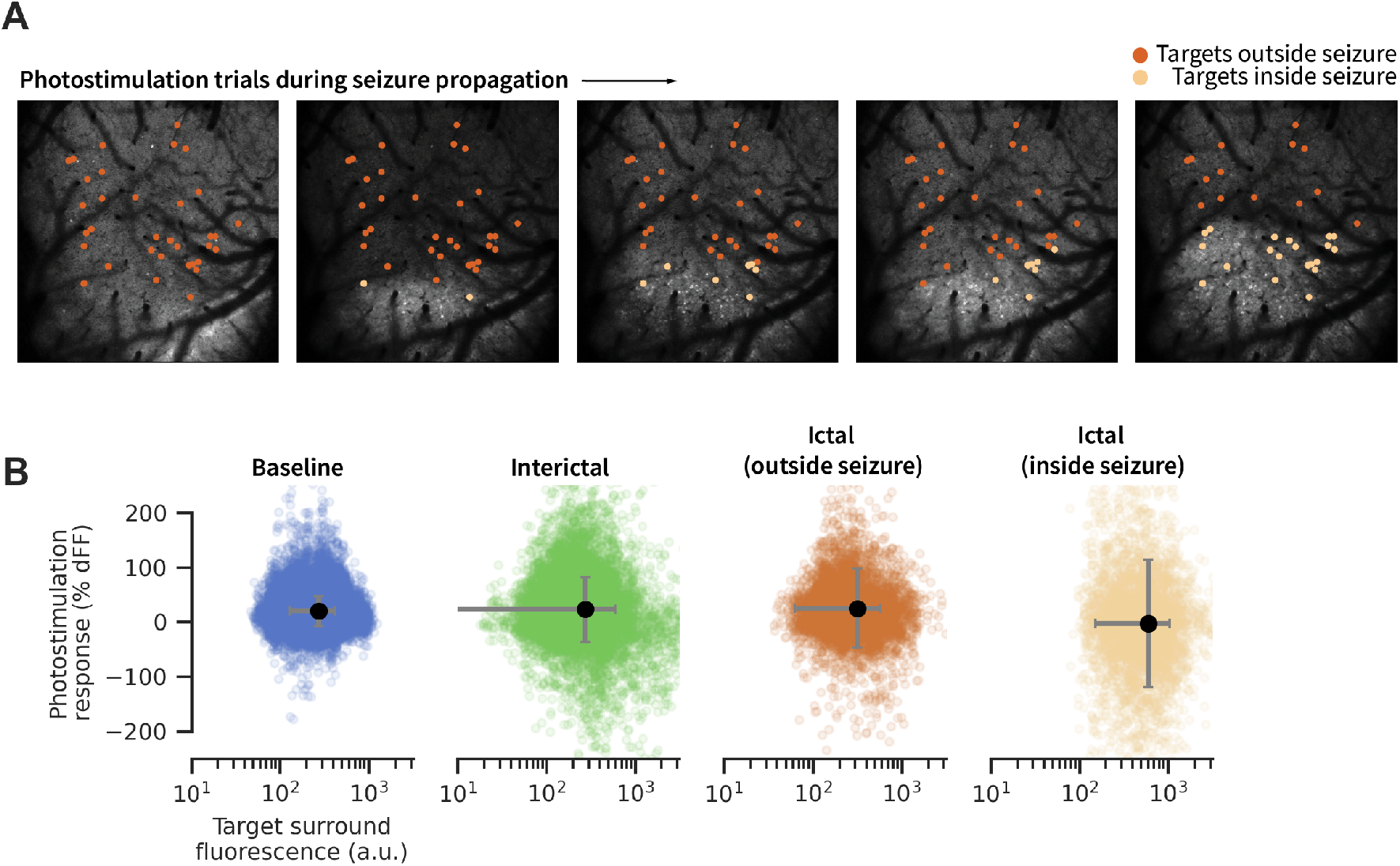
Seizure boundary classification A. Calcium fluorescence images of consecutive photostimulation trials during seizure propagation of a single representative ictal event from one experiment. Fluorescence images are 3 secs average around time of photostimulation trial. All photostimulation targets in each experiment were individually classified as being inside or outside the seizure wavefront boundary on a given photostimulation trial during ictal events. Photostimulation trials with unclear seizure wavefront locations (i.e. wavefront has not yet reached FOV, or wavefront has passed FOV) did not receive this classification for targets’ location. B. Relationship of surrounding fluorescence (raw) signal to photostimulation response of all targets and photostimulation trials across all experiments. Ictal photostimulation trials were further split by individual targets’ location inside/outside the seizure wavefront boundary at each individual photostimulation trial.

## Notes

### Competing Interest Statement

The authors have declared no competing interest.

## References

1. Carl C. H. Petersen. Whole-Cell Recording of Neuronal Membrane Potential during Behavior. Neuron, 95(6):1266–1281, September 2017. ISSN 0896-6273. doi: 10.1016/j.neuron.2017.06.049.

2. Ingrid E. Scheffer, Samuel Berkovic, Giuseppe Capovilla, Mary B. Connolly, Jacqueline French, Laura Guilhoto, Edouard Hirsch, Satish Jain, Gary W. Mathern, Solomon L. Moshé, Douglas R. Nordli, Emilio Perucca, Torbjörn Tomson, Samuel Wiebe, Yue-Hua Zhang, and Sameer M. Zuberi. ILAE classification of the epilepsies: Position paper of the ILAE Commission for Classification and Terminology. Epilepsia, 58(4):512–521, 2017. ISSN 1528-1167. doi: 10.1111/epi.13709. _eprint: https://onlinelibrary.wiley.com/doi/pdf/10.1111/epi.13709.

3. Simon Shorvon, Renzo Guerrini, Steven Schachter, and Eugen Trinka, editors. The Causes of Epilepsy: Common and Uncommon Causes in Adults and Children. Cambridge University Press, Cambridge, 2 edition, 2019. ISBN 978-1-108-42075-4. doi: 10.1017/9781108355209.

4. William W. Lytton. Computer modelling of epilepsy. Nature Reviews Neuroscience, 9(8): 626–637, August 2008. ISSN 1471-003X, 1471-0048. doi: 10.1038/nrn2416.

5. Samuel Wiebe and Nathalie Jette. Pharmacoresistance and the role of surgery in difficult to treat epilepsy. Nature Reviews Neurology, 8(12):669–677, December 2012. ISSN 1759-4766. doi: 10.1038/nrneurol.2012.181. Number: 12 Publisher: Nature Publishing Group.

6. Catherine A. Schevon, Shennan A. Weiss, Guy McKhann, Robert R. Goodman, Rafael Yuste, Ronald G. Emerson, and Andrew J. Trevelyan. Evidence of an inhibitory restraint of seizure activity in humans. Nature Communications, 3(1):1060, January 2012. ISSN 2041-1723. doi: 10.1038/ncomms2056.

7. Elliot H. Smith, Jyun-you Liou, Tyler S. Davis, Edward M. Merricks, Spencer S. Kellis, Shennan A. Weiss, Bradley Greger, Paul A. House, Guy M. McKhann Ii, Robert R. Goodman, Ronald G. Emerson, Lisa M. Bateman, Andrew J. Trevelyan, and Catherine A. Schevon. The ictal wavefront is the spatiotemporal source of discharges during spontaneous human seizures. Nature Communications, 7(1):11098, September 2016. ISSN 2041-1723. doi: 10.1038/ncomms11098.

8. D. A. Prince and B. J. Wilder. Control mechanisms in cortical epileptogenic foci. “Surround” inhibition. Archives of Neurology, 16(2):194–202, February 1967. ISSN 0003-9942. doi: 10.1001/archneur.1967.00470200082007.

9. Theodore H. Schwartz and Tobias Bonhoeffer. In vivo optical mapping of epileptic foci and surround inhibition in ferret cerebral cortex. Nature Medicine, 7(9):1063–1067, September 2001. ISSN 1546-170X. doi: 10.1038/nm0901-1063. Number: 9 Publisher: Nature Publishing Group.

10. Jyun-you Liou, Hongtao Ma, Michael Wenzel, Mingrui Zhao, Eliza Baird-Daniel, Elliot H Smith, Andy Daniel, Ronald Emerson, Rafael Yuste, Theodore H Schwartz, and Catherine A Schevon. Role of inhibitory control in modulating focal seizure spread. Brain, 141(7):2083– 2097, July 2018. ISSN 0006-8950, 1460-2156. doi: 10.1093/brain/awy116.

11. R. Ryley Parrish, Neela K. Codadu, Connie Mackenzie-Gray Scott, and Andrew J. Trevelyan. Feedforward inhibition ahead of ictal wavefronts is provided by both parvalbumin- and somatostatin-expressing interneurons. The Journal of Physiology, 597(8):2297–2314, April 2019. ISSN 0022-3751. doi: 10.1113/JP277749.

12. Matthew Weston. Jumping to Conclusions about Focal Seizure Spread. Epilepsy Currents, 18(6):394–395, November 2018. ISSN 1535-7597, 1535-7511. doi: 10.5698/1535-7597.18.6.394.

13. Mario Cammarota, Gabriele Losi, Angela Chiavegato, Micaela Zonta, and Giorgio Carmignoto. Fast spiking interneuron control of seizure propagation in a cortical slice model of focal epilepsy: Fast-spiking interneurons in focal seizure propagation. The Journal of Physiology, 591(4):807–822, February 2013. ISSN 00223751. doi: 10.1113/jphysiol.2012.238154.

14. M. Sessolo, I. Marcon, S. Bovetti, G. Losi, M. Cammarota, G. M. Ratto, T. Fellin, and G. Carmignoto. Parvalbumin-Positive Inhibitory Interneurons Oppose Propagation But Favor Generation of Focal Epileptiform Activity. Journal of Neuroscience, 35(26):9544–9557, July 2015. ISSN 0270-6474, 1529-2401. doi: 10.1523/JNEUROSCI.5117-14.2015.

15. Andrew J. Trevelyan and Catherine A. Schevon. How inhibition influences seizure propagation. Neuropharmacology, 69:45–54, June 2013. ISSN 0028-3908. doi: 10.1016/j.neuropharm.2012.06.015.

16. A. J. Trevelyan, D. Sussillo, B. O. Watson, and R. Yuste. Modular Propagation of Epileptiform Activity: Evidence for an Inhibitory Veto in Neocortex. Journal of Neuroscience, 26(48): 12447–12455, November 2006. ISSN 0270-6474, 1529-2401. doi: 10.1523/JNEUROSCI.2787-06.2006.

17. Jyun-you Liou, Elliot H Smith, Lisa M Bateman, Samuel L Bruce, Guy M McKhann, Robert R Goodman, Ronald G Emerson, Catherine A Schevon, and Lf Abbott. A model for focal seizure onset, propagation, evolution, and progression. eLife, 9:e50927, March 2020. ISSN 2050-084X. doi: 10.7554/eLife.50927.

18. Tahra L. Eissa, Koen Dijkstra, Christoph Brune, Ronald G. Emerson, Michel J. A. M. van Putten, Robert R. Goodman, Guy M. McKhann, Catherine A. Schevon, Wim van Drongelen, and Stephan A. van Gils. Cross-scale effects of neural interactions during human neocortical seizure activity. Proceedings of the National Academy of Sciences, 114(40): 10761–10766, October 2017. doi: 10.1073/pnas.1702490114. Publisher: Proceedings of the National Academy of Sciences.

19. H. Alfonsa, E. M. Merricks, N. K. Codadu, M. O. Cunningham, K. Deisseroth, C. Racca, and A. J. Trevelyan. The Contribution of Raised Intraneuronal Chloride to Epileptic Network Activity. Journal of Neuroscience, 35(20):7715–7726, May 2015. ISSN 0270-6474, 1529-2401. doi: 10.1523/JNEUROSCI.4105-14.2015.

20. Vincent Magloire, Jonathan Cornford, Andreas Lieb, Dimitri M. Kullmann, and Ivan Pavlov. KCC2 overexpression prevents the paradoxical seizure-promoting action of somatic inhibition. Nature Communications, 10(1):1225, December 2019. ISSN 2041-1723. doi: 10.1038/s41467-019-08933-4.

21. Michael Chang, Joshua A. Dian, Suzie Dufour, Lihua Wang, Homeira Moradi Chameh, Meera Ramani, Liang Zhang, Peter L. Carlen, Thilo Womelsdorf, and Taufik A. Valiante. Brief activation of GABAergic interneurons initiates the transition to ictal events through post-inhibitory rebound excitation. Neurobiology of Disease, 109:102–116, January 2018. ISSN 09699961. doi: 10.1016/j.nbd.2017.10.007.

22. Esther Krook-Magnuson, Caren Armstrong, Mikko Oijala, and Ivan Soltesz. On-demand optogenetic control of spontaneous seizures in temporal lobe epilepsy. Nature Communications, 4(1):1376, June 2013. ISSN 2041-1723. doi: 10.1038/ncomms2376.

23. Latefa Yekhlef, Gian Luca Breschi, Laura Lagostena, Giovanni Russo, and Stefano Taverna. Selective activation of parvalbumin- or somatostatin-expressing interneurons triggers epileptic seizurelike activity in mouse medial entorhinal cortex. Journal of Neurophysiology, 113(5):1616–1630, March 2015. ISSN 0022-3077. doi: 10.1152/jn.00841.2014. Publisher: American Physiological Society.

24. Kenneth D. Harris and Thomas D. Mrsic-Flogel. Cortical connectivity and sensory coding. Nature, 503(7474):51–58, November 2013. ISSN 0028-0836, 1476-4687. doi: 10.1038/nature12654.

25. Julie A. Harris, Stefan Mihalas, Karla E. Hirokawa, Jennifer D. Whitesell, Hannah Choi, Amy Bernard, Phillip Bohn, Shiella Caldejon, Linzy Casal, Andrew Cho, Aaron Feiner, David Feng, Nathalie Gaudreault, Charles R. Gerfen, Nile Graddis, Peter A. Groblewski, Alex M. Henry, Anh Ho, Robert Howard, Joseph E. Knox, Leonard Kuan, Xiuli Kuang, Jerome Lecoq, Phil Lesnar, Yaoyao Li, Jennifer Luviano, Stephen McConoughey, Marty T. Mortrud, Maitham Naeemi, Lydia Ng, Seung Wook Oh, Benjamin Ouellette, Elise Shen, Staci A. Sorensen, Wayne Wakeman, Quanxin Wang, Yun Wang, Ali Williford, John W. Phillips, Allan R. Jones, Christof Koch, and Hongkui Zeng. Hierarchical organization of cortical and thalamic connectivity. Nature, 575(7781):195–202, November 2019. ISSN 1476-4687. doi: 10.1038/s41586-019-1716-z. Number: 7781 Publisher: Nature Publishing Group.

26. Kenneth D Harris and Gordon M G Shepherd. The neocortical circuit: themes and variations. Nature Neuroscience, 18(2):170–181, February 2015. ISSN 1097-6256, 1546-1726. doi: 10.1038/nn.3917.

27. Hal Blumenfeld. What Is a Seizure Network? Long-Range Network Consequences of Focal Seizures. In Helen E. Scharfman and Paul S. Buckmaster, editors, Issues in Clinical Epileptology: A View from the Bench, Advances in Experimental Medicine and Biology, pages 63–70. Springer Netherlands, Dordrecht, 2014. ISBN 978-94-017-8914-1. doi: 10.1007/978-94-017-8914-1_5.

28. Richard J. Burman and R. Ryley Parrish. The Widespread Network Effects of Focal Epilepsy. Journal of Neuroscience, 38(38):8107–8109, September 2018. ISSN 0270-6474, 1529-2401. doi: 10.1523/JNEUROSCI.1471-18.2018. Publisher: Society for Neuroscience Section: Journal Club.

29. ManKin Choy, Ehsan Dadgar-Kiani, Greg O. Cron, Ben A. Duffy, Florian Schmid, Bradley J. Edelman, Mazen Asaad, Russell W. Chan, Shahabeddin Vahdat, and Jin Hyung Lee. Repeated hippocampal seizures lead to brain-wide reorganization of circuits and seizure propagation pathways. Neuron, 110(2):221–236.e4, January 2022. ISSN 08966273. doi: 10.1016/j.neuron.2021.10.010.

30. Jeanne T Paz, Thomas J Davidson, Eric S Frechette, Bruno Delord, Isabel Parada, Kathy Peng, Karl Deisseroth, and John R Huguenard. Closed-loop optogenetic control of thalamus as a tool for interrupting seizures after cortical injury. Nature Neuroscience, 16(1):64–70, January 2013. ISSN 1097-6256, 1546-1726. doi: 10.1038/nn.3269.

31. Laurent Sheybani, Gwenaël Birot, Alessandro Contestabile, Margitta Seeck, Jozsef Zoltan Kiss, Karl Schaller, Christoph M. Michel, and Charles Quairiaux. Electrophysiological Evidence for the Development of a Self-Sustained Large-Scale Epileptic Network in the Kainate Mouse Model of Temporal Lobe Epilepsy. Journal of Neuroscience, 38(15):3776–3791, April 2018. ISSN 0270-6474, 1529-2401. doi: 10.1523/JNEUROSCI.2193-17.2018.

32. Yi Wang, Cenglin Xu, Zhenghao Xu, Caihong Ji, Jiao Liang, Ying Wang, Bin Chen, Xiaohua Wu, Feng Gao, Shuang Wang, Yi Guo, Xiaoming Li, Jianhong Luo, Shumin Duan, and Zhong Chen. Depolarized GABAergic Signaling in Subicular Microcircuits Mediates Generalized Seizure in Temporal Lobe Epilepsy. Neuron, 95(1):92–105.e5, July 2017. ISSN 08966273. doi: 10.1016/j.neuron.2017.06.004.

33. Joshua E. Motelow, Wei Li, Qiong Zhan, Asht M. Mishra, Robert N.S. Sachdev, Geoffrey Liu, Abhijeet Gummadavelli, Zaina Zayyad, Hyun Seung Lee, Victoria Chu, John P. Andrews, Dario J. Englot, Peter Herman, Basavaraju G. Sanganahalli, Fahmeed Hyder, and Hal Blumenfeld. Decreased Subcortical Cholinergic Arousal in Focal Seizures. Neuron, 85 (3):561–572, February 2015. ISSN 08966273. doi: 10.1016/j.neuron.2014.12.058.

34. Michael A Long and Albert K Lee. Intracellular recording in behaving animals. Current Opinion in Neurobiology, 22(1):34–44, February 2012. ISSN 0959-4388. doi: 10.1016/j.conb.2011.10.013.

35. Lyle J. Borg-Graham, Cyril Monier, and Yves Frégnac. Visual input evokes transient and strong shunting inhibition in visual cortical neurons. Nature, 393(6683):369–373, May 1998. ISSN 1476-4687. doi: 10.1038/30735. Number: 6683 Publisher: Nature Publishing Group.

36. Alain Destexhe, Michael Rudolph, and Denis Paré. The high-conductance state of neocortical neurons in vivo. Nature Reviews Neuroscience, 4(9):739–751, September 2003. ISSN 1471-003X, 1471-0048. doi: 10.1038/nrn1198.

37. Denis Paré, Eric Shink, Hélène Gaudreau, Alain Destexhe, and Eric J. Lang. Impact of Spontaneous Synaptic Activity on the Resting Properties of Cat Neocortical Pyramidal Neurons In Vivo. Journal of Neurophysiology, 79(3):1450–1460, March 1998. ISSN 0022-3077, 1522-1598. doi: 10.1152/jn.1998.79.3.1450.

38. Daniel F. English, Adrien Peyrache, Eran Stark, Lisa Roux, Daniela Vallentin, Michael A. Long, and György Buzsáki. Excitation and Inhibition Compete to Control Spiking during Hippocampal Ripples: Intracellular Study in Behaving Mice. Journal of Neuroscience, 34(49): 16509–16517, December 2014. ISSN 0270-6474, 1529-2401. doi: 10.1523/JNEUROSCI.2600-14.2014. Publisher: Society for Neuroscience Section: Articles.

39. Adam M Packer, Lloyd E Russell, Henry W P Dalgleish, and Michael Häusser. Simultaneous all-optical manipulation and recording of neural circuit activity with cellular resolution in vivo. Nature Methods, 12(2):140–146, February 2015. ISSN 1548-7091, 1548-7105. doi: 10.1038/nmeth.3217.

40. John Peter Rickgauer, Karl Deisseroth, and David W Tank. Simultaneous cellular-resolution optical perturbation and imaging of place cell firing fields. Nature Neuroscience, 17(12): 1816–1824, December 2014. ISSN 1097-6256, 1546-1726. doi: 10.1038/nn.3866.

41. Alberto Morales-Villagrán and Ricardo Tapia. Preferential stimulation of glutamate release by 4-aminopyridine in rat striatum in vivo. Neurochemistry International, 28(1):35–40, January 1996. ISSN 0197-0186. doi: 10.1016/0197-0186(95)00064-F.

42. Michael Wenzel, Jordan P. Hamm, Darcy S. Peterka, and Rafael Yuste. Acute Focal Seizures Start As Local Synchronizations of Neuronal Ensembles. The Journal of Neuroscience, 39(43):8562–8575, October 2019. ISSN 0270-6474, 1529-2401. doi: 10.1523/JNEUROSCI.3176-18.2019.

43. Michael Wenzel, Jordan P. Hamm, Darcy S. Peterka, and Rafael Yuste. Reliable and Elastic Propagation of Cortical Seizures In Vivo. Cell Reports, 19(13):2681–2693, June 2017. ISSN 22111247. doi: 10.1016/j.celrep.2017.05.090.

44. L. Federico Rossi, Robert C. Wykes, Dimitri M. Kullmann, and Matteo Carandini. Focal cortical seizures start as standing waves and propagate respecting homotopic connectivity. Nature Communications, 8(1):217, December 2017. ISSN 2041-1723. doi: 10.1038/s41467-017-00159-6.

45. György Buzsáki, Costas A. Anastassiou, and Christof Koch. The origin of extracellular fields and currents — EEG, ECoG, LFP and spikes. Nature Reviews Neuroscience, 13(6):407– 420, June 2012. ISSN 1471-003X, 1471-0048. doi: 10.1038/nrn3241.

46. Alex A. Legaria, Bridget A. Matikainen-Ankney, Ben Yang, Biafra Ahanonu, Julia A. Licholai, Jones G. Parker, and Alexxai V. Kravitz. Fiber photometry in striatum reflects primarily non-somatic changes in calcium. Nature Neuroscience, 25(9):1124–1128, September 2022. ISSN 1546-1726. doi: 10.1038/s41593-022-01152-z. Number: 9 Publisher: Nature Publishing Group.

47. Jeffry S. Isaacson and Massimo Scanziani. How Inhibition Shapes Cortical Activity. Neuron, 72(2):231–243, October 2011. ISSN 08966273. doi: 10.1016/j.neuron.2011.09.027.

48. Alfonso Renart, Jaime de la Rocha, Peter Bartho, Liad Hollender, Néstor Parga, Alex Reyes, and Kenneth D. Harris. The Asynchronous State in Cortical Circuits. Science, 327(5965): 587–590, January 2010. ISSN 0036-8075, 1095-9203. doi: 10.1126/science.1179850.

49. Robert T. Graham, R. Ryley Parrish, Laura Alberio, Emily L. Johnson, Laura J. Owens, and Andrew J. Trevelyan. Synergistic positive feedback underlying seizure initiation. preprint, Neuroscience, March 2021.

50. Lloyd E. Russell, Henry W. P. Dalgleish, Rebecca Nutbrown, Oliver M. Gauld, Dustin Herrmann, Mehmet Fişek, Adam M. Packer, and Michael Häusser. All-optical interrogation of neural circuits in behaving mice. Nature Protocols, pages 1–42, April 2022. ISSN 1750-2799. doi: 10.1038/s41596-022-00691-w. Publisher: Nature Publishing Group.

51. Naiyan Chen, Hiroki Sugihara, and Mriganka Sur. An acetylcholine-activated microcircuit drives temporal dynamics of cortical activity. Nature Neuroscience, 18(6):892–902, June 2015. ISSN 1097-6256, 1546-1726. doi: 10.1038/nn.4002.

52. Simon Musall, Matthew T. Kaufman, Ashley L. Juavinett, Steven Gluf, and Anne K. Churchland. Single-trial neural dynamics are dominated by richly varied movements. Nature Neuroscience, 22(10):1677–1686, October 2019. ISSN 1546-1726. doi: 10.1038/s41593-019-0502-4. Number: 10 Publisher: Nature Publishing Group.

53. Carsen Stringer, Marius Pachitariu, Nicholas Steinmetz, Matteo Carandini, and Kenneth D. Harris. High-dimensional geometry of population responses in visual cortex. Nature, 571 (7765):361–365, July 2019. ISSN 0028-0836, 1476-4687. doi: 10.1038/s41586-019-1346-5.

54. Carsen Stringer, Marius Pachitariu, Nicholas Steinmetz, Charu Bai Reddy, Matteo Carandini, and Kenneth D. Harris. Spontaneous behaviors drive multidimensional, brainwide activity. Science, 364(6437):eaav7893, April 2019. ISSN 0036-8075, 1095-9203. doi: 10.1126/science.aav7893.

55. C. van Vreeswijk and H. Sompolinsky. Chaotic Balanced State in a Model of Cortical Circuits. Neural Computation, 10(6):1321–1371, August 1998. ISSN 0899-7667, 1530-888X. doi: 10.1162/089976698300017214.

56. Pariya Salami, Mia Borzello, Mark A. Kramer, M. Brandon Westover, and Sydney S. Cash. Quantification of seizure termination patterns reveals limited pathways to seizure end. Technical report, medRxiv, medRxiv, March 2021. Type: article.

57. Norman K. So and Warren T. Blume. The postictal EEG. Epilepsy & Behavior: E&B, 19(2): 121–126, October 2010. ISSN 1525-5069. doi: 10.1016/j.yebeh.2010.06.033.

58. James F. A. Poulet and Carl C. H. Petersen. Internal brain state regulates membrane potential synchrony in barrel cortex of behaving mice. Nature, 454(7206):881–885, August 2008. ISSN 0028-0836, 1476-4687. doi: 10.1038/nature07150.

59. Selmaan N. Chettih and Christopher D. Harvey. Single-neuron perturbations reveal feature-specific competition in V1. Nature, 567(7748):334–340, March 2019. ISSN 0028-0836, 1476-4687. doi: 10.1038/s41586-019-0997-6.

60. Henry WP Dalgleish, Lloyd E Russell, Adam M Packer, Arnd Roth, Oliver M Gauld, Francesca Greenstreet, Emmett J Thompson, and Michael Häusser. How many neurons are sufficient for perception of cortical activity? eLife, 9:e58889, October 2020. ISSN 2050-084X. doi: 10.7554/eLife.58889.

61. Alan R. Mardinly, Ian Antón Oldenburg, Nicolas C. Pégard, Savitha Sridharan, Evan H. Lyall, Kirill Chesnov, Stephen G. Brohawn, Laura Waller, and Hillel Adesnik. Precise multimodal optical control of neural ensemble activity. Nature Neuroscience, 21(6):881–893, June 2018. ISSN 1097-6256, 1546-1726. doi: 10.1038/s41593-018-0139-8.

62. Ian Antón Oldenburg, William D. Hendricks, Gregory Handy, Kiarash Shamardani, Hayley A. Bounds, Brent Doiron, and Hillel Adesnik. The logic of recurrent circuits in the primary visual cortex. preprint, Neuroscience, September 2022.

63. Lloyd E. Russell, Zidan Yang, Pei Lynn Tan, Mehmet Fişek, Adam M. Packer, Henry W.P. Dalgleish, Selmaan Chettih, Christopher D. Harvey, and Michael Häusser. The influence of visual cortex on perception is modulated by behavioural state. preprint, Neuroscience, July 2019.

64. Elodie Fino, Adam M. Packer, and Rafael Yuste. The logic of inhibitory connectivity in the neocortex. The Neuroscientist: A Review Journal Bringing Neurobiology, Neurology and Psychiatry, 19(3):228–237, June 2013. ISSN 1089-4098. doi: 10.1177/1073858412456743.

65. Carl Holmgren, Tibor Harkany, Björn Svennenfors, and Yuri Zilberter. Pyramidal cell communication within local networks in layer 2/3 of rat neocortex. The Journal of Physiology, 551(Pt 1):139–153, August 2003. ISSN 0022-3751. doi: 10.1113/jphysiol.2003.044784.

66. Christoph Kapfer, Lindsey L. Glickfeld, Bassam V. Atallah, and Massimo Scanziani. Supralinear increase of recurrent inhibition during sparse activity in the somatosensory cortex. Nature Neuroscience, 10(6):743–753, June 2007. ISSN 1546-1726. doi: 10.1038/nn1909. Number: 6 Publisher: Nature Publishing Group.

67. A. J. Trevelyan, D. Sussillo, and R. Yuste. Feedforward Inhibition Contributes to the Control of Epileptiform Propagation Speed. Journal of Neuroscience, 27(13):3383–3387, March 2007. ISSN 0270-6474, 1529-2401. doi: 10.1523/JNEUROSCI.0145-07.2007.

68. Manuel Valero, Ipshita Zutshi, Euisik Yoon, and György Buzsáki. Probing subthreshold dynamics of hippocampal neurons by pulsed optogenetics. Science, 375(6580):570–574, February 2022. ISSN 0036-8075, 1095-9203. doi: 10.1126/science.abm1891.

69. Luis Carrillo-Reid, Shuting Han, Weijian Yang, Alejandro Akrouh, and Rafael Yuste. Controlling Visually Guided Behavior by Holographic Recalling of Cortical Ensembles. Cell, 178 (2):447–457.e5, July 2019. ISSN 00928674. doi: 10.1016/j.cell.2019.05.045.

70. James H. Marshel, Yoon Seok Kim, Timothy A. Machado, Sean Quirin, Brandon Benson, Jonathan Kadmon, Cephra Raja, Adelaida Chibukhchyan, Charu Ramakrishnan, Masatoshi Inoue, Janelle C. Shane, Douglas J. McKnight, Susumu Yoshizawa, Hideaki E. Kato, Surya Ganguli, and Karl Deisseroth. Cortical layer–specific critical dynamics triggering perception. Science, 365(6453):eaaw5202, August 2019. ISSN 0036-8075, 1095-9203. doi: 10.1126/science.aaw5202.

71. James M. Rowland, Thijs L. van der Plas, Matthias Loidolt, Robert M. Lees, Joshua Keeling, Jonas Dehning, Thomas Akam, Viola Priesemann, and Adam M. Packer. Perception and propagation of activity through the cortical hierarchy is determined by neural variability. preprint, Neuroscience, bioRxiv, December 2021.

72. Prisca R. Bauer, Roland D. Thijs, Robert J. Lamberts, Demetrios N. Velis, Gerhard H. Visser, Else A. Tolner, Josemir W. Sander, Fernando H. Lopes da Silva, and Stiliyan N. Kalitzin. Dynamics of convulsive seizure termination and postictal generalized EEG suppression. Brain, 140(3):655–668, March 2017. ISSN 0006-8950. doi: 10.1093/brain/aww322.

73. Eric Halgren, Thomas L. Babb, and Paul H. Crandall. Post-EEG Seizure Depression of Human Limbic Neurons Is Not Détérmined by Their Response to Probable Hypoxia. Epilepsia, 18(1):89–93, 1977. ISSN 1528-1167. doi: 10.1111/j.1528-1157.1977.tb05590.x. _eprint: https://onlinelibrary.wiley.com/doi/pdf/10.1111/j.1528-1157.1977.tb05590.x.

74. Fred A. Lado and Solomon L. Moshé. How do seizures stop? Epilepsia, 49(10): 1651–1664, 2008. ISSN 1528-1167. doi: 10.1111/j.1528-1167.2008.01669.x. _eprint: https://onlinelibrary.wiley.com/doi/pdf/10.1111/j.1528-1167.2008.01669.x.

75. Frédéric Zubler, Andreas Steimer, Heidemarie Gast, and Kaspar A. Schindler. Seizure termination. International Review of Neurobiology, 114:187–207, 2014. ISSN 2162-5514. doi: 10.1016/B978-0-12-418693-4.00008-X.

76. Frances S Chance, L. F Abbott, and Alex D Reyes. Gain Modulation from Background Synaptic Input. Neuron, 35(4):773–782, August 2002. ISSN 0896-6273. doi: 10.1016/S0896-6273(02)00820-6.

77. C. Koch, R. Douglas, and U. Wehmeier. Visibility of synaptically induced conductance changes: theory and simulations of anatomically characterized cortical pyramidal cells. Journal of Neuroscience, 10(6):1728–1744, June 1990. ISSN 0270-6474, 1529-2401. doi: 10.1523/JNEUROSCI.10-06-01728.1990. Publisher: Society for Neuroscience Section: Articles.

78. Marlene Bartos, Imre Vida, and Peter Jonas. Synaptic mechanisms of synchronized gamma oscillations in inhibitory interneuron networks. Nature Reviews Neuroscience, 8(1):45–56, January 2007. ISSN 1471-003X, 1471-0048. doi: 10.1038/nrn2044.

79. Joseph V. Raimondo, Louise Kay, Tommas J. Ellender, and Colin J. Akerman. Optogenetic silencing strategies differ in their effects on inhibitory synaptic transmission. Nature Neuroscience, 15(8):1102–1104, August 2012. ISSN 1546-1726. doi: 10.1038/nn.3143. Number: 8 Publisher: Nature Publishing Group.

80. Joseph V Raimondo, Blake A Richards, and Melanie A Woodin. Neuronal chloride and excitability — the big impact of small changes. Current Opinion in Neurobiology, 43:35–42, April 2017. ISSN 0959-4388. doi: 10.1016/j.conb.2016.11.012.

81. Luke Campagnola, Stephanie C. Seeman, Thomas Chartrand, Lisa Kim, Alex Hoggarth, Clare Gamlin, Shinya Ito, Jessica Trinh, Pasha Davoudian, Cristina Radaelli, Mean-Hwan Kim, Travis Hage, Thomas Braun, Lauren Alfiler, Julia Andrade, Phillip Bohn, Rachel Dalley, Alex Henry, Sara Kebede, Mukora Alice, David Sandman, Grace Williams, Rachael Larsen, Corinne Teeter, Tanya L. Daigle, Kyla Berry, Nadia Dotson, Rachel Enstrom, Melissa Gorham, Madie Hupp, Samuel Dingman Lee, Kiet Ngo, Philip R. Nicovich, Lydia Potekhina, Shea Ransford, Amanda Gary, Jeff Goldy, Delissa McMillen, Trangthanh Pham, Michael Tieu, La’Akea Siverts, Miranda Walker, Colin Farrell, Martin Schroedter, Cliff Slaughterbeck, Charles Cobb, Richard Ellenbogen, Ryder P. Gwinn, C. Dirk Keene, Andrew L. Ko, Jeffrey G. Ojemann, Daniel L. Silbergeld, Daniel Carey, Tamara Casper, Kirsten Crichton, Michael Clark, Nick Dee, Lauren Ellingwood, Jessica Gloe, Matthew Kroll, Josef Sulc, Herman Tung, Katherine Wadhwani, Krissy Brouner, Tom Egdorf, Michelle Maxwell, Medea McGraw, Christina Alice Pom, Augustin Ruiz, Jasmine Bomben, David Feng, Nika Hejazinia, Shu Shi, Aaron Szafer, Wayne Wakeman, John Phillips, Amy Bernard, Luke Esposito, Florence D. D’Orazi, Susan Sunkin, Kimberly Smith, Bosiljka Tasic, Anton Arkhipov, Staci Sorensen, Ed Lein, Christof Koch, Gabe Murphy, Hongkui Zeng, and Tim Jarsky. Local connectivity and synaptic dynamics in mouse and human neocortex. Science, 375(6585): eabj5861, March 2022. ISSN 0036-8075, 1095-9203. doi: 10.1126/science.abj5861.

82. A. M. Packer and R. Yuste. Dense, Unspecific Connectivity of Neocortical Parvalbumin-Positive Interneurons: A Canonical Microcircuit for Inhibition? Journal of Neuroscience, 31(37):13260–13271, September 2011. ISSN 0270-6474, 1529-2401. doi: 10.1523/JNEUROSCI.3131-11.2011.

83. R W Gerchberg and W O Saxton. A Practical Algorithm for the Determination of Phase from Image and Diffraction Plane Pictures. Optik, 35:237–246, 1972.

84. Carsen Stringer, Tim Wang, Michalis Michaelos, and Marius Pachitariu. Cellpose: a generalist algorithm for cellular segmentation. Nature Methods, 18(1):100–106, January 2021. ISSN 1548-7105. doi: 10.1038/s41592-020-01018-x. Number: 1 Publisher: Nature Publishing Group.

